# Deciphering the reciprocal regulation of the Pyk2–Src activation complex

**DOI:** 10.64898/2026.07.20.739604

**Authors:** Alexzandrea Woudenberg, Kallie M. Zuidema, Tania M. Palhano Zanela, Karen G. Romero Bello, Eric S. Underbakke

## Abstract

Pyk2 and Src are non-receptor tyrosine kinases that assemble into a complex to promote mutual activation. The Pyk2 FERM domain mediates autoinhibition by direct interaction with the kinase. Src autoinhibition is maintained by intramolecular interactions of the SH3 and SH2 domains with internal ligands, including an inhibitory C-terminal phosphotyrosine. FERM disengagement permits Pyk2 autophosphorylation of the FERM–kinase linker to generate a scaffolding site for Src recruitment. Although Src-mediated phosphorylation of the Pyk2 activation loop is well-established, the mechanism by which complex formation activates Src remains unclear. We reconstituted defined phosphorylation and regulatory states of both kinases, combining site-directed mutagenesis with phosphosite-resolved activity profiling to dissect the reciprocal regulation. Src phosphorylates the Pyk2 activation loop via an ordered, self-primed dual phosphorylation mechanism. Although activated Pyk2 productively phosphorylates the Src activation loop, Pyk2–Src complex formation does not significantly increase activation loop phosphorylation above the rate of Src autophosphorylation alone. Conversely, autoinhibited Src strictly requires Pyk2 scaffolding engagement to productively phosphorylate Pyk2. Pyk2 therefore relieves Src autoinhibition by presenting competing scaffolding sites to disengage intramolecular SH3 and SH2 conformational constraints. The results distinguish activation loop phosphorylation from the conformational competence required for substrate phosphorylation. We propose that autophosphorylated Pyk2 functions principally as a conformational activator and scaffolding platform, opening Src for phosphatase-mediated removal of the inhibitory C-terminal phosphosite and committing both Pyk2 and Src to downstream substrate phosphorylation.

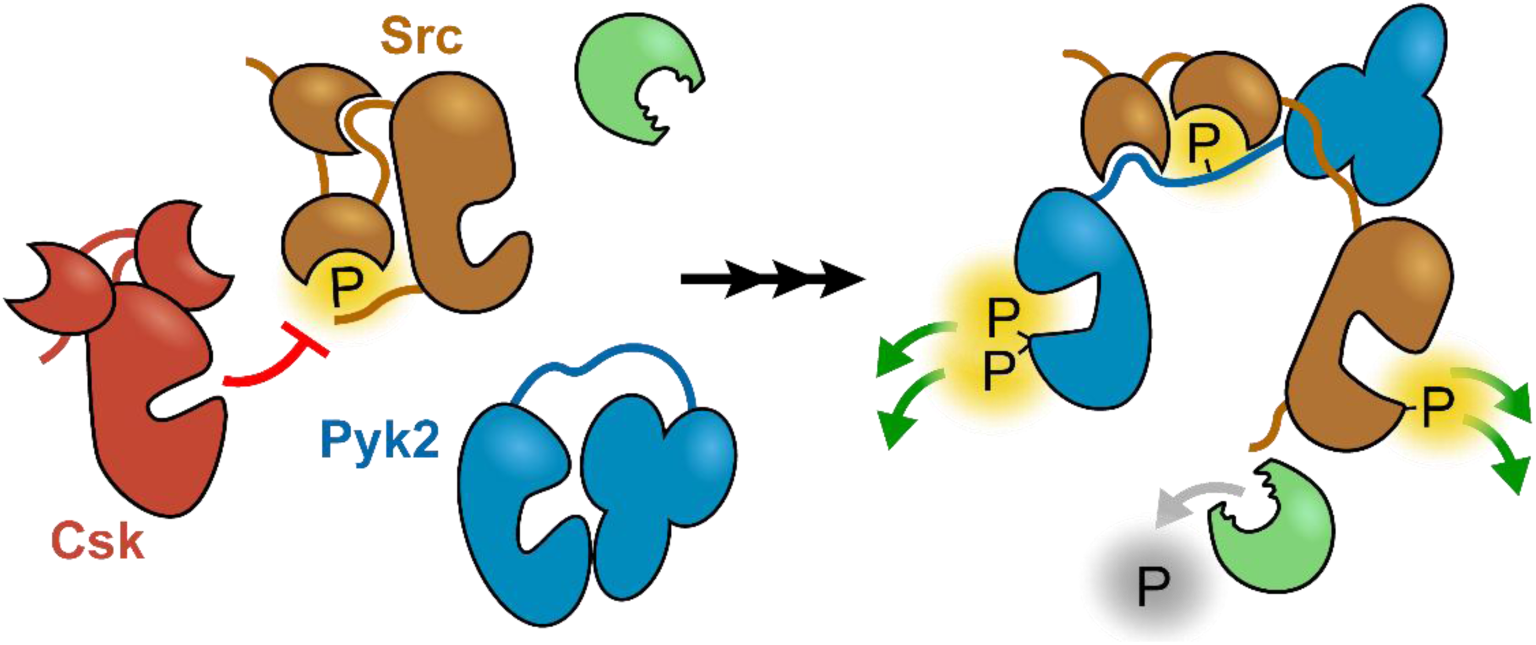

## Introduction

Proline-rich tyrosine kinase 2 (Pyk2) and Src are non-receptor tyrosine kinases that collaborate to promote mutual activation and signaling (1, 2). In cellular signaling pathways, strict regulation of kinase activity is essential for maintaining responsivity, robustness, and fidelity. Kinases transition between multiple activity states, modulated by phosphorylation, localization, and regulatory interactions (3). In particular, phosphorylation of the kinase activation loop organizes the active site, enhancing substrate accessibility and promoting higher phosphotransfer activity (4). Activation of kinases through conformational changes and phosphorylation requires coordinated protein–protein interactions to ensure robust signal transduction.

Pyk2 is the sole paralog of focal adhesion kinase (FAK), exhibiting 65% sequence similarity and comparable domain organization (5, 6). Despite shared evolutionary origins, the two kinases have diverged in expression patterns, subcellular localization, activation mechanisms, and signaling outputs (6–10). FAK is localized to the membrane and activated by lipid interactions and clustering at focal adhesions (11–13) with signaling roles in cell adhesion, migration, proliferation, and embryonic development (14, 15). Conversely, Pyk2 retains a cytosolic localization and responds to Ca²⁺ flux in neuronal cells (6, 16, 17). In other cell types such as endothelial cells and megakaryocytes, Pyk2 preserves more traditional FAK-like localization and functions (7, 18). Neuronal Pyk2 is enriched in the post synaptic density and is associated with synaptic plasticity (19, 20). Pyk2 dysfunction is implicated in neurological disorders, such as Huntington’s and Alzheimer’s disease (1, 20, 21).

Despite a notable divergence in localization and activation stimuli, Pyk2 retains key signatures of the FAK multistage activation mechanism, including the requirement of Src family kinase (SFK) recruitment to achieve activation loop phosphorylation (22, 23). Both FAK and Pyk2 contain an N-terminal regulatory FERM (protein 4.1/ezrin/radixin/moesin) domain, a central tyrosine kinase, and a C-terminal FAT (focal adhesion targeting) domain (**Fig. 1A**). Pyk2 and FAK are inhibited via FERM interaction with the kinase, blocking access to the active site (**Fig. 1B**) (24, 25). While the details remain ambiguous, a combination of clustering and FERM rearrangements relieve FERM inhibition (12, 17, 26). Disinhibition of FAK and Pyk2 is characterized by the opening of the multidomain architecture, freeing the kinase for limited phosphotransfer function (12, 17, 27). Subsequently, the FERM–kinase linker tyrosine, Y402 (Y397 in FAK), undergoes autophosphorylation (22, 23, 28). The autophosphorylated linker tyrosine (pY402) and a proximal proline-rich region serve as a high affinity docking site for the tandem Src SH2 and SH3 domains (**Fig. 1A**) (29). Src then phosphorylates Pyk2 activation loop residues Y579/Y580 (Y576/Y577 in FAK), transitioning the kinase to a fully active state (22, 23). Accordingly, FAK and Pyk2 serve dual roles as both signaling enzymes and scaffolds (30). For example, the scaffolding interaction of Src with FAK/Pyk2 provides a platform for activated Src to target pathway-specific downstream effectors (31, 32).

**Figure 1.**
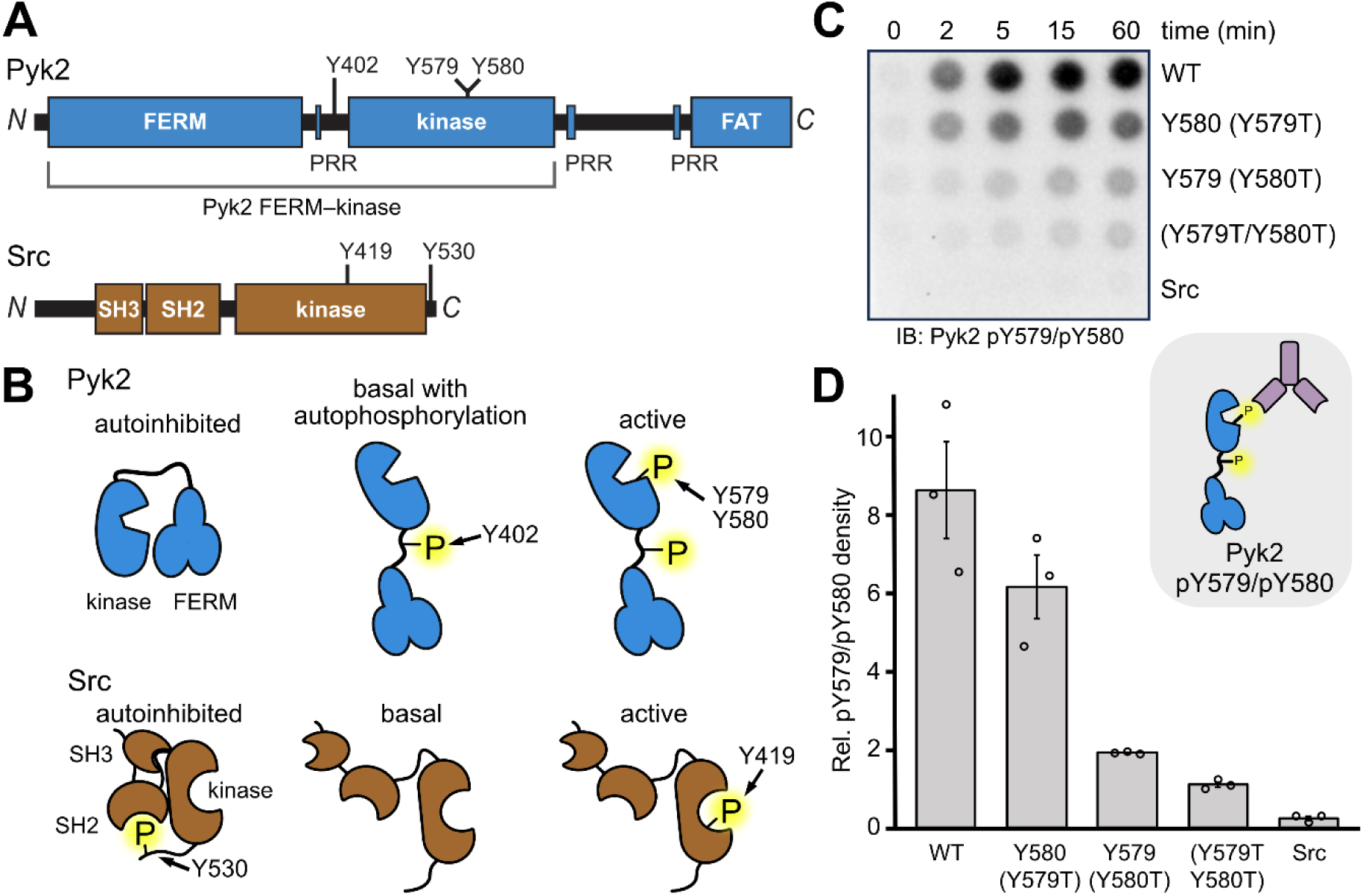
Src phosphorylation preference of Pyk2 activation loop residues. (A) Pyk2 and Src domain organizations scaled to sequence length with phosphotyrosine sites and proline-rich regions (PRR) annotated. The Pyk2 FERM–kinase construct (residues 20–692) is also labeled. (B) Cartoon schematics of Pyk2 (top) and Src (bottom) phosphorylation sites and conformations governing regulatory states. (C) Phosphorylation time courses of Pyk2 FERM–kinase variants (0.5 μM) in the presence of Src (0.2 μM). Pyk2 WT, Y579T, Y580T, and Y579T/Y580T were pre-incubated with ATP at 37 °C to fully phosphorylate Y402. Tyrosine phosphorylation of the Pyk2 activation loop was detected via dot blotting using Pyk2 pY579/pY580-specific primary antibody. (D) Pyk2 activation loop variants demonstrate a preference for Src phosphorylation. Quantification of activation loop phosphorylation (pY579/pY580) replicates (Fig. S1) from independent reactions (n = 3) at 5 min. Error bars represent standard error. *Inset:* Key to immunoblotting antibody recognition site in C and D.

Src family kinases are well-known signaling hubs involved in numerous cellular pathways (33, 34). SFK functions include cell proliferation, motility, cell cycle regulation, and receptor-mediated signal transduction (35, 36). Src is the prototypical member of the Src kinase family and the first discovered tyrosine kinase proto-oncogene (34). Src dysfunction is implicated in various types of cancer, establishing Src as a key therapeutic target (37, 38).

Src contains an N-terminal SH3 domain followed by an SH2 domain and a C-terminal tyrosine kinase (**Fig. 1A**). The conformational arrangement of Src arises from an interplay of phosphorylation and intramolecular interactions of the SH2 and SH3 domains (34, 39). Conformational rearrangement underlies SFK regulation, where competition for SH2/SH3 interactions is linked to kinase activation. Src autoinhibition is maintained by intramolecular interactions that wedge the kinase domain into a catalytically incompetent conformation (40, 41). Phosphorylation of the C-terminus (Y530) by CSK (C-terminal Src kinase) generates an internal ligand for the SH2 domain, while the SH3 domain binds a pseudo-ligand in the SH2–kinase linker, forming suboptimal intramolecular interactions (**Fig. 1B**) (40–44). Src activation is achieved through displacement of the intramolecular SH2/SH3 interactions via binding to competitive ligands, activation loop phosphorylation, and concomitant dephosphorylation of the C-terminal Y530 by co-localized phosphatases (34, 45). For example, receptor tyrosine kinases (e.g., epidermal growth factor receptor and platelet-derived growth factor receptor) present intracellular motifs to recruit and activate Src (46, 47). Displacement of the SH– or SH3-mediated intramolecular interactions instigates a scaffolding-based activation (41, 48). Phosphorylation of the Src activation loop (Y419) stabilizes the active conformation of the kinase domain to confer the highest levels of phosphotransfer activity (**Fig. 1B**) (45, 49).

In cells, formation of the Pyk2–Src or FAK–Src complex leads to activation of not only Pyk2/FAK, but Src as well (2, 28, 50). Overexpression of Pyk2, but not Pyk2 Y402F, leads to increased Src activation loop phosphorylation (pY419) (2). Both FAK and Pyk2 linker autophosphorylation sites have been found to bind the Src SH2 domain (2, 28, 51). Src-mediated Pyk2 activation loop phosphorylation is more significantly impaired when access to pY402 is blocked compared to the nearby proline-rich region (52). Taken together, the FAK and Pyk2 autophosphorylation sites serve as productive Src SH2 ligands, and complex formation increases Src activity. Src activation likely relies on release of autoinhibition when Pyk2 presents tandem SH2/SH3 docking sites to outcompete Src intramolecular interactions. Nevertheless, the progression of interactions and phosphorylations leading to mutual Src and Pyk2 phosphorylation remains unclear.

In cells, the interaction between Pyk2 and Src ultimately provokes activation of both kinases, but the mechanistic details remain unclear. Src may require pre-activation by upstream signaling factors prior to Pyk2 engagement. Alternatively, Src may remain autoinhibited until Pyk2 binding relieves conformational inhibition. Nevertheless, it is still unknown whether Pyk2 is responsible for Src activation loop phosphorylation or if scaffolding alone is sufficient to promote Src autophosphorylation. Here, we aim to distinguish direct phosphorylation from conformational regulation and clarify the mechanistic basis of mutual kinase activation and cooperative signaling roles.

## Results

### Src phosphorylation of Pyk2 activation loop

Co-incubation of Src and Pyk2 *in vitro* can drive both kinases to maximal phosphorylation and activity (53), a process that presumably involves phosphorylation of both Src and Pyk2 activation loops. Interestingly, Pyk2 is among several kinases that require multiple activation loop tyrosine phosphorylations, including FAK, IRK, MET, and Jak2 (23, 54–56). The phosphorylated activation loop tyrosines stabilize the Pyk2 active site via a network of electrostatic interactions with the surrounding basic pocket (53). Pyk2, in either the basal or active state, exhibits negligible autophosphorylation of either activation loop residue, Y579 or Y580 (**Fig. S1**). Intact protein mass spectrometry corroborates the role of Src in Pky2 activation loop phosphorylation (**Fig. S2**). Prior to ATP incubation, Pyk2 exhibits a single unphosphorylated mass. Following autophosphorylation, the signal shifts to a mass consistent with Pyk2 monophosphorylation (+80 Da) at Y402. Addition of Src quickly yields a single species with two additional phosphorylations. As such, Src is seemingly responsible for phosphorylation of both adjacent residues. However, upon monophosphorylation of either tyrosine, the local electrostatics and sterics of the Pyk2 activation loop are significantly altered, necessitating flexible substrate recognition by Src. If Src is responsible for phosphorylation of both tyrosines in the Pyk2 activation loop, we wondered whether Src exhibits ordered or preferred phosphorylation of either Pyk2 residue.

To investigate whether the Src-mediated dual phosphorylation favored either activation loop residue, Y579 or Y580, we generated non-phosphorylatable Pyk2 activation loop variants Y579T, Y580T, and Y579T/Y580T. Src readily monophosphorylates Y580 (Y579T) on a similar timescale as WT. However, Y579 (Y580T) phosphorylation is reduced to the background levels exhibited by the non-phosphorylatable Y579T/Y580T variant (**Fig. 1C and 1D, S1**). Therefore, Src preferentially phosphorylates residue Y580. Evidently, the altered electrostatics of monophosphorylation or conformational rearrangement of the activation loop primes subsequent Y579 phosphorylation.

### Pyk2 phosphorylation of Src

In cells, Pyk2 and FAK interact with Src, promoting Src activation (2, 22, 50). However, it is not clear whether Pyk2 is directly responsible for Src activation loop (Y419) phosphorylation. Given the scaffolded proximity of the activation loops of both proteins during Src phosphorylation of Pyk2, we sought to test whether Pyk2 can reciprocally phosphorylate Src Y419 *in vitro*. To focus on Pyk2-mediated phosphorylation of the Src activation loop, we restricted Src activity by employing a kinase dead (K298M) Src variant. Activation loop phosphorylation of K298M Src was tested with either basal (pY402, unphosphorylated loop) or fully active (pY579/pY580) Pyk2. While basal Pyk2 cannot phosphorylate Src Y419 (**Fig. 2A**), active Pyk2 robustly phosphorylates Src Y419 (**Fig. 2B**). The extent of Pyk2-mediated phosphorylation of Src Y419 is ∼60% of the level observed for WT Src autophosphorylation (**Fig. 2C, S3**). Notably, Src activity may be inflated relative to Pyk2, as Src autophosphorylation is subject to autocatalytic amplification as the reaction progresses (i.e., the product, pY419 Src, is activated for rapid, continued autophosphorylation in trans).

**Figure 2.**
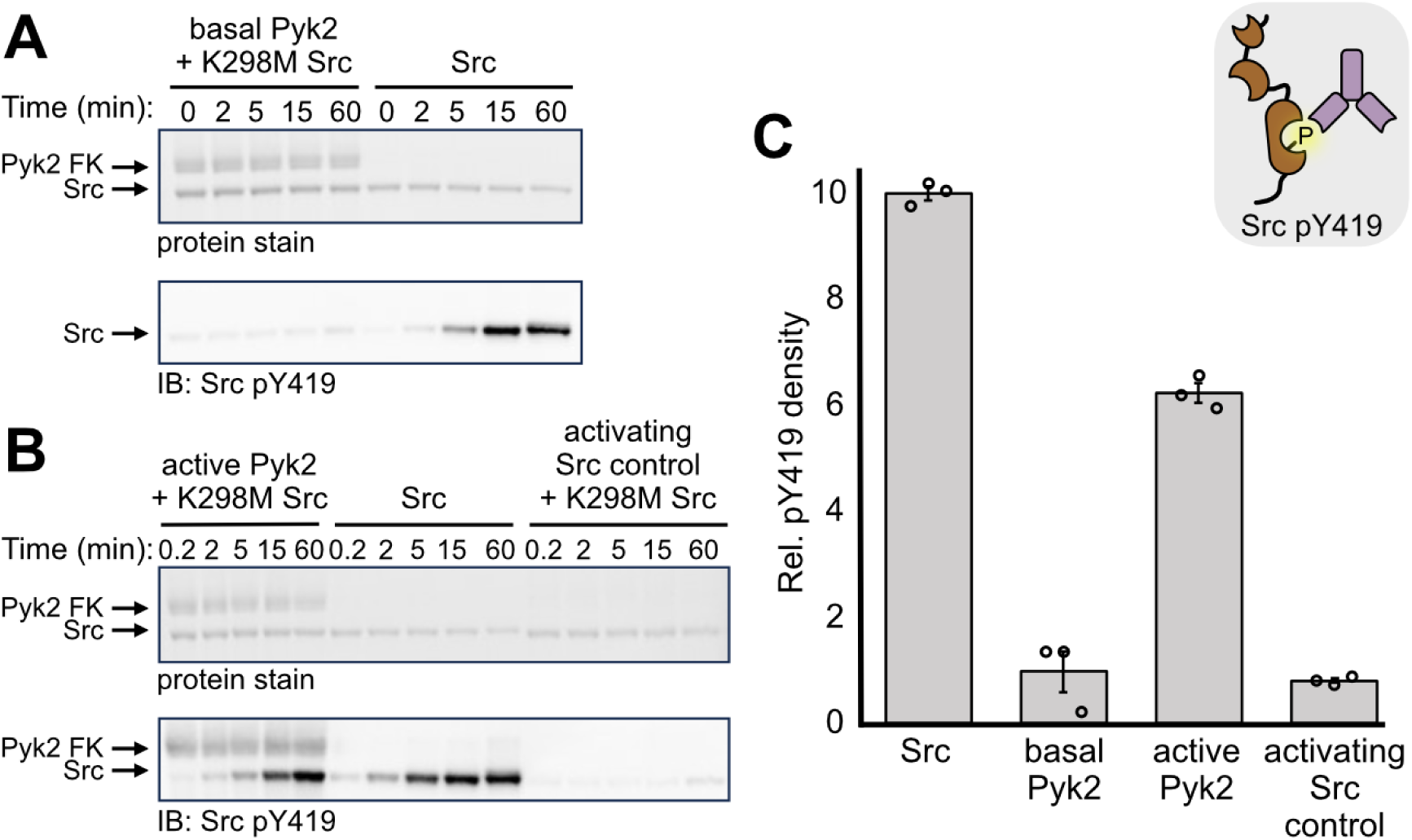
Pyk2 phosphorylation of Src activation loop. (A) Comparison of Src activation loop phosphorylation mediated by basal Pyk2 vs. Src autophosphorylation. Pyk2 FERM–kinase (1 μM) was pre-incubated with ATP for 15 min to generate pY402 Pyk2, and subsequently, K298M Src (1 μM) was added to the reaction (left). WT Src (1 μM) autophosphorylation was monitored in the absence of Pyk2 (right). Phosphorylation of the Src activation loop was assessed via Western blotting using Src pY419-specific primary antibody. (B) Comparison of Src activation loop phosphorylation mediated by fully phosphorylated, active Pyk2 FERM–kinase vs. Src autophosphorylation. Active, fully phosphorylated Pyk2 FERM–kinase was generated via pre-incubation with minimal Src (115:1 stoichiometric ratio of Pyk2 to Src). Specifically, Pyk2 (23 μM) was pre-incubated with Src (0.2 μM) and ATP for 180 min. Active pY579/pY580 Pyk2 (1 μM) was incubated with equimolar K298M Src (left). For comparison, WT Src (1 μM) autophosphorylation was assessed without Pyk2 (center). To control for the background of residual Src used to activate Pyk2, a mock reaction was performed omitting Pyk2 FERM–kinase with the same overall dilution of catalytic Src (right). Phosphorylation of the Src activation loop was assessed by Western blotting using Src pY419-specific primary antibody. (C) Phosphorylation of Src Y419 was quantified by densitometry from replicates (**Fig. S3A**) of independent reactions (n = 3) at 15 min. *Inset:* Key to immunoblotting antibody recognition site in A–C.

### Src activation loop phosphorylation is not required to activate Pyk2

Src activation loop phosphorylation serves two primary roles: enhancing catalytic activity and influencing substrate specificity (45, 57, 58). We established that Pyk2 can promote Src activation loop phosphorylation, however Pyk2 activation loop phosphorylation is a prerequisite (**Fig. 2**), an apparent circular dependency. If Pyk2 contributes to Src activation loop phosphorylation in cells, an upstream kinase is needed for prior Pyk2 activation loop phosphorylation. We wondered whether unphosphorylated Src basal activity sufficed to target and phosphorylate the Pyk2 activation loop after recruitment to the phosphorylated Pyk2 linker site, pY402.

To assess whether Src activation loop phosphorylation is a required for the efficient recognition and phosphorylation of the Pyk2 activation loop, we generated a non-phosphorylatable loop variant, Y419F Src. We then challenged WT Src or Y419F Src to phosphorylate the pY402 Pyk2 FERM-kinase activation loop *in vitro*. Notably, both WT and Y419F Src readily phosphorylate Pyk2 residues Y579/Y580, demonstrating that basal Src activity is sufficient to phosphorylate the Pyk2 activation loop (**Fig. 3A**).

**Figure 3.**
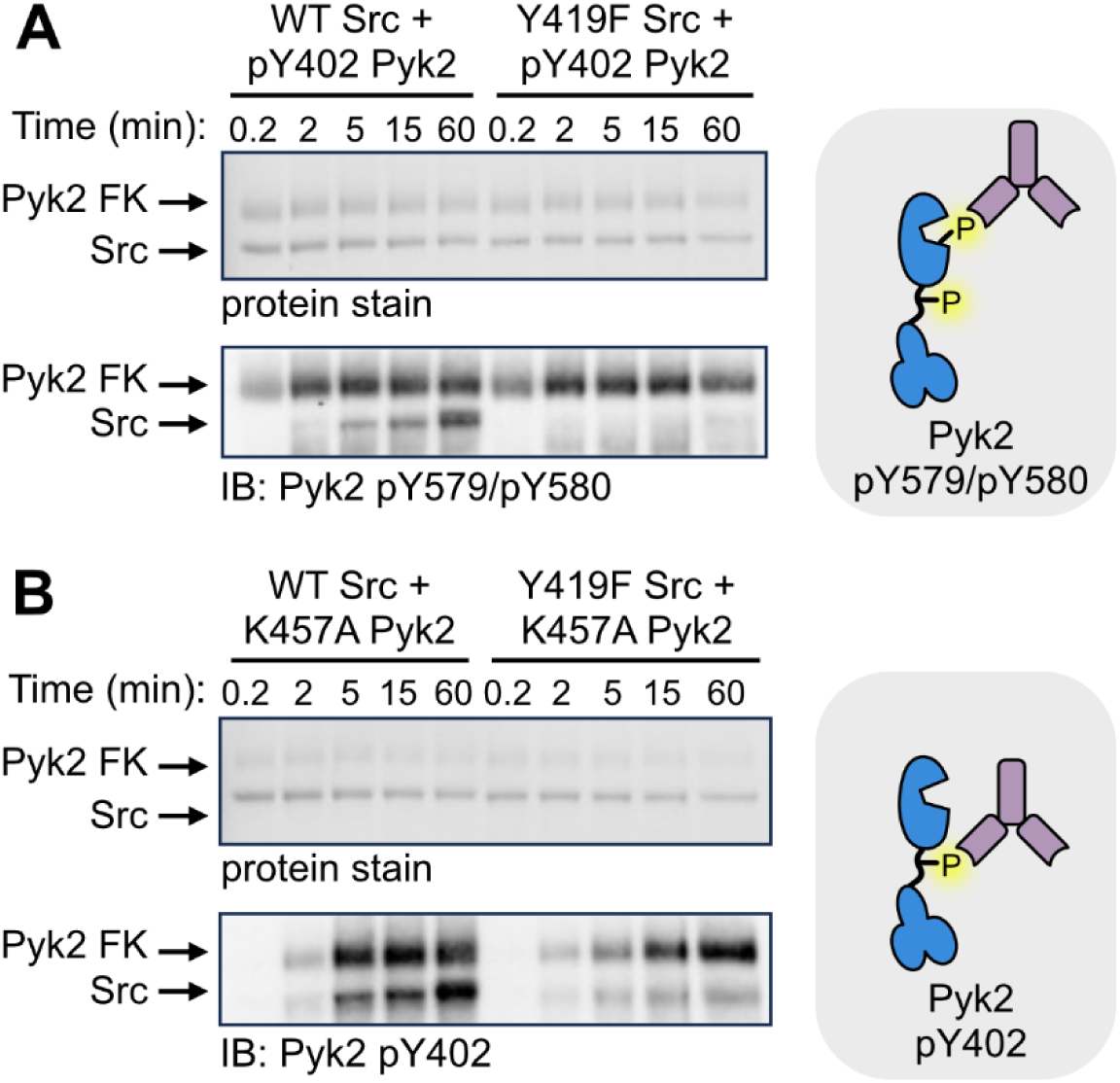
Src does not require activation loop phosphorylation to phosphorylate Pyk2. (A) Phosphorylation time course of pY402 Pyk2 FERM–kinase (0.5 μM) with WT or Y419F Src (0.5 μM). Pyk2 FERM–kinase was pre-incubated with ATP for 15 min at 37 °C to autophosphorylate the linker (pY402 Pyk2). Src-mediated phosphorylation of the Pyk2 activation loop was monitored via Western blotting using Pyk2 pY579/pY580-specific primary antibody. (B) Phosphorylation time course of kinase dead K457A Pyk2 FERM–kinase (0.5 μM) with WT or Y419F Src (0.5 μM). Tyrosine phosphorylation of the Pyk2 autophosphorylation site was detected via Western blotting using Pyk2 pY402-specific primary antibody. *Insets:* Keys to immunoblotting antibody recognition sites in A and B, respectively.

While Src can prime phosphorylation of the Pyk2 FERM—kinase linker site (Y402) in some contexts (59), trans autophosphorylation promoted by clustering is thought to be the primary mechanism governing Pyk2 linker phosphorylation (17, 22, 53). Nevertheless, prior investigations of Src-mediated Pyk2 linker phosphorylation were complicated by concomitant activation loop phosphorylation and autocatalytic amplification of Pyk2 activity. To clarify the extent to which Src can phosphorylate Pyk2 site Y402, Y419F Src was tested with a kinase dead Pyk2 variant, K457A. Intriguingly, the Y419F variant can also phosphorylate the Y402 autophosphorylation site, albeit less rapidly than WT Src (**Fig. 3B**).

### Pyk2 scaffolding outcompetes Src autoinhibition

While Pyk2-mediated phosphorylation (pY419) can promote Src activation (**Fig. 2B and 2C**), we also wanted to test whether Pyk2 can play a role in conformational stimulation of Src activity. Several signaling systems trigger Src family kinase activation by outcompeting the autoinhibitory intramolecular SH2/SH3 interactions (**Fig. 1B**) (42, 47, 48, 60). Pyk2 and FAK are known to require Src SH2 binding for activation loop phosphorylation, indicating that the phosphorylated FERM—kinase linkers serve as preferred Src docking sites (2, 29, 52). Indeed, the tandem SH2/SH3 ligands presented by the phosphorylated Pyk2 FERM—kinase linker (i.e., pY402 and proline-rich region) offer a potentially high affinity Src scaffolding site. Therefore, autophosphorylated Pyk2 may exert conformational control over Src activity by outcompeting autoinhibitory SH2/SH3 interactions, independent of activation loop phosphorylation. We aimed to test the impact of Pyk2 scaffolding on Src activity using defined kinase regulatory states *in vitro*.

To determine if Pyk2 can serve as a conformational activator of Src, we first needed to generate autoinhibited Src. Upon autophosphorylation of the Src activation loop, the kinase domain is locked in an active state (45, 49). To focus on conformational control and rule out Src autophosphorylation, we started with non-phosphorylatable Y419F Src. Autoinhibited Y419F Src was generated by treatment with CSK, the kinase responsible for phosphorylation of the Src C-terminus (Y530) to form the intramolecular SH2 ligand (**Fig. S5**). To confirm that CSK-treated pY530 Src activity is inhibited, we deployed Y402F Pyk2 as a non-scaffolding phosphoacceptor substrate. The Pyk2 pY402 site is the primary determinant of Src docking (2, 28, 51, 52), therefore Y402F Pyk2 presents known Src phosphorylation target sites (Y579/Y580) without perturbing the Src conformation. Indeed, basal activity Y419F Src (no CSK treatment) still achieves Pyk2 activation loop phosphorylation (**Fig. 4A**). However, the CSK-treated, autoinhibited Y419F Src exhibits no detectable Y402F Pyk2 phosphorylation (**Fig. 4A**). Therefore, our *in vitro* system recapitulates strict Src autoinhibition upon CSK-mediated C-terminal phosphorylation.

**Figure 4.**
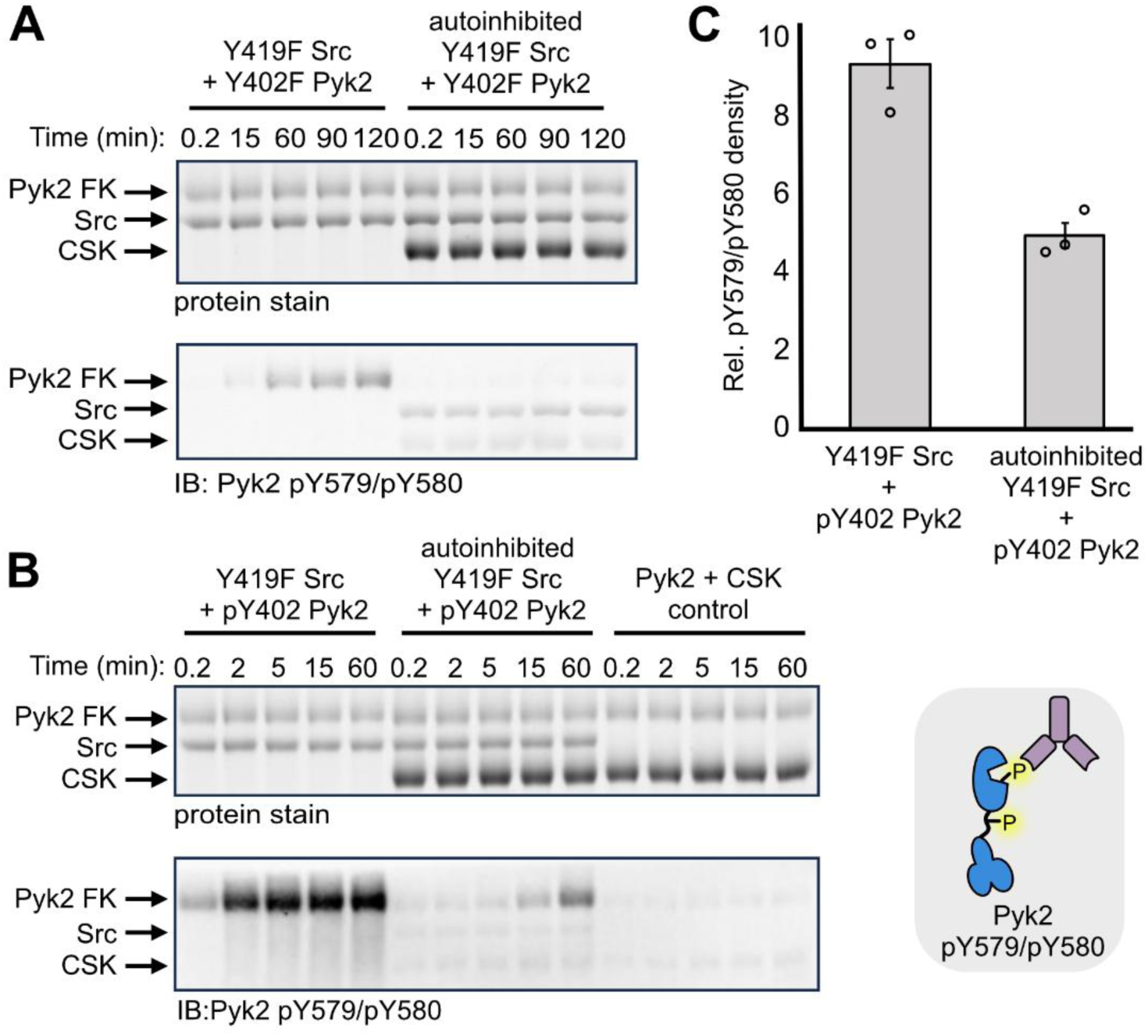
Pyk2 pY402 scaffolding outcompetes Y419F Src autoinhibition. (A) Kinase activity time course of basal (unphosphorylated) or autoinhibited (pY530) Y419F Src (2 μM) with Y402F Pyk2 FERM–kinase (2 μM). Autoinhibited Y419F Src (2 μM) was a product of incubation with 5 μM CSK and ATP for 10 min. Phosphorylation of the Pyk2 activation loop was monitored via Western blotting using Pyk2 pY579/pY580-specific primary antibody. (B) Kinase activity time course of open or autoinhibited Y419F Src (2 μM) with pY402 Pyk2 FERM–kinase (2 μM). Pyk2 was incubated with ATP for 15 min at 37 °C to generate pY402 Pyk2. Basal or autoinhibited (pY530) Y419F Src was combined with pY402 Pyk2 (2 μM), and Src kinase activity was monitored via Western blotting using Pyk2 pY579/pY580-specific primary antibody. (C) Activation loop phosphorylation of pY402 Pyk2 was quantified by densitometry from independent reactions (n = 3) at 60 min with open or autoinhibited Y419F Src (**Fig. S4A**). *Inset:* Key to immunoblotting antibody recognition sites in A–C.

Next, we directly tested whether Pyk2 with autophosphorylated FERM—kinase linker (pY402) is sufficient to stimulate autoinhibited (pY530) Y419F Src activity. Either open basal (no CSK treatment) or autoinhibited (pY530) Y419F Src was incubated with pY402 Pyk2. To reveal whether Pyk2 can outcompete Src autoinhibition, we monitored Pyk2 activation loop phosphorylation as an indicator of Src activation. A control with Pyk2 and CSK alone exhibited no activation loop phosphorylation, confirming CSK specificity for Src (**Fig. 4B**). As previously observed (**Fig. 3A**), basal state Y419F Src readily phosphorylates the Pyk2 activation loop (**Fig. 4B**). While pY530 constrains Src to negligible activity with non-scaffolding Pyk2 substrate (**Fig. 4A**), pY530 autoinhibition is overcome and activity restored when Src is presented with pY402 Pyk2 as substrate (**Fig. 4B**). Indeed, pY402 Pyk2 is sufficient to stimulate pY530 Src activity to ∼50% of the uninhibited Src basal activity (**Fig. 4C, S4**). Therefore, Pyk2 linker phosphorylation can relieve Src autoinhibition, independent of Src activation loop phosphorylation.

Thus far, we observed that fully phosphorylated, active (pY579/pY580) Pyk2 readily phosphorylates the Src activation loop (**Fig. 2B-C),** while Src does not require activation loop phosphorylation to activate Pyk2 (**Fig. 3A**). These findings support a regulatory model involving a reciprocal kinase phosphorylation cascade. In cells, both native kinases access dynamic conformational and phosphorylation-induced regulatory landscapes. Autocatalytic amplification can rapidly propagate activation loop phosphorylation via trans autophosphorylation, while scaffolding can simultaneously promote conformational rearrangement and substrate proximity. To test whether an ordered, reciprocal phosphorylation mechanism is feasible, we reconstituted a defined Pyk2/Src system *in vitro* using WT kinases with full phosphorylation potential. By balancing CSK and tyrosine phosphatase YopH activities, we generated WT Src and Pyk2 FERM–kinase in defined starting states of (de)phosphorylation.

Briefly, Src was autoinhibited as previously described by incubating with CSK and ATP to phosphorylate the C-terminus. Autoinhibited pY530 Src was then treated with YopH to reset the Src activation loop to the dephosphorylated state (**Fig. S5**). The autoinhibited Src conformation is known to protect pY530 from phosphatase activity (61), as we confirmed by immunoblotting (**Fig. S5**). YopH was subsequently inhibited with orthovandate to allow for subsequent kinase assays. In tandem, immunoblotting was performed to confirm that pY402 Pyk2 was generated via autophosphorylation with no detectable activation loop phosphorylation (**Fig. S6**).

To test whether CSK/phosphatase treatment recapitulated autoinhibition in WT pY530 Src, we compared the activities of CSK/phosphatase-treated WT Src with open state (untreated) WT Src using non-scaffolding, Y402F Pyk2 as a phosphoacceptor substrate (**Fig. 5A**). Similar to Y419F Src (**Fig. 4A**), autoinhibited WT Src exhibits no Y402F Pyk2 activation loop phosphorylation (**Fig. 5A**). Open (unphosphorylated) Src robustly phosphorylates pY402 Pyk2 (**Fig. 5B-C**), also mirroring Y419F Src (**Fig. 4B-C**). Presented with the potential for scaffolding interactions, autoinhibited WT Src achieves significant phosphorylation of the pY402 Pyk2 activation loop (**Fig. 5C**), albeit with a significant lag phase early in the time course (**Fig. 5B**).

**Figure 5.**
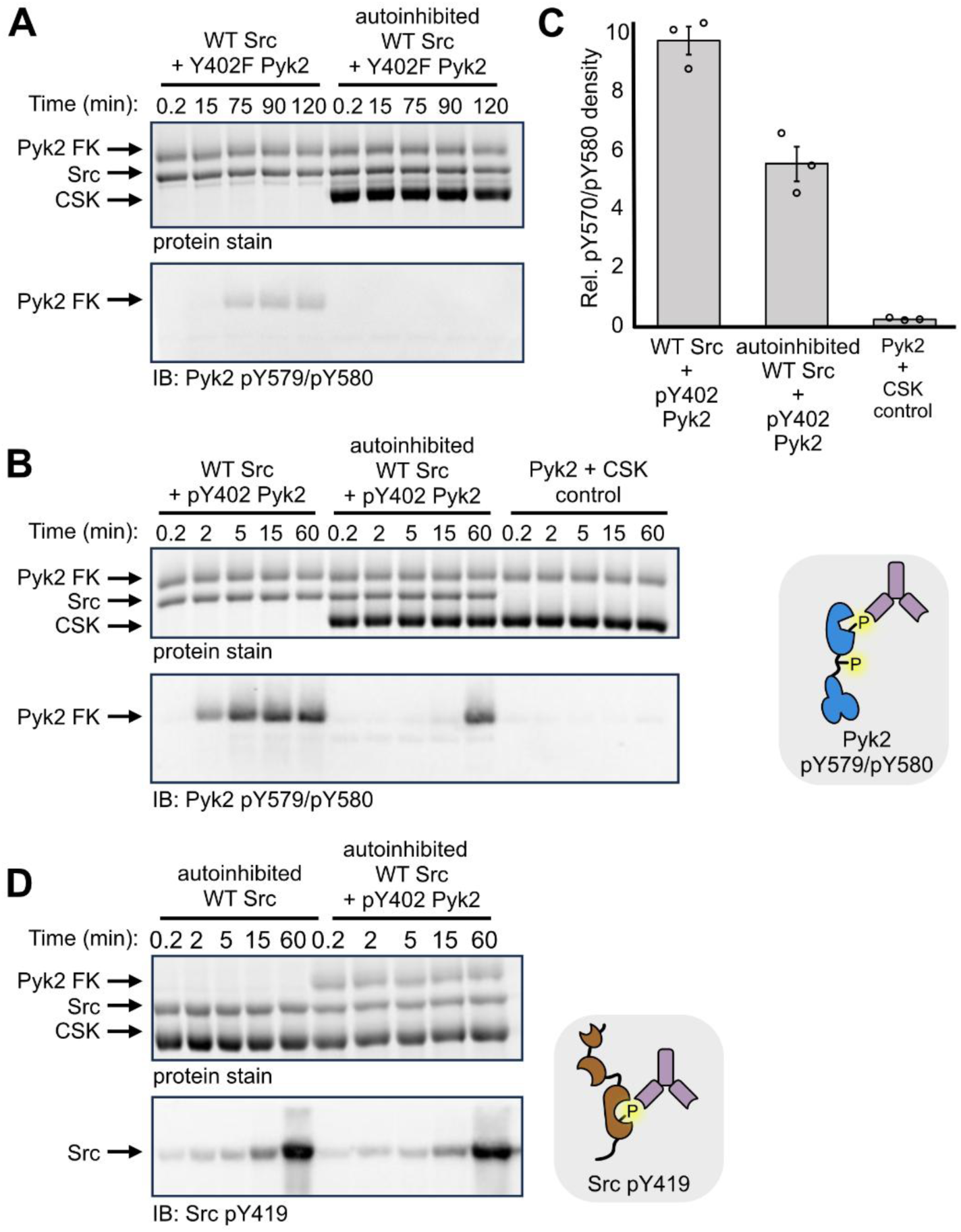
Reconstitution of reciprocal activation loop phosphorylation of Pyk2 and Src. (A) Kinase activity time course of open (initially unphosphorylated) or autoinhibited (pY530) WT Src (2 μM) with Y402F Pyk2 FERM–kinase (2 μM). Phosphorylation of the Pyk2 activation loop was detected via Western blotting using Pyk2 pY579/pY580-specific primary antibody. (B) Kinase activity time course of open or autoinhibited WT Src (2 μM) with WT Pyk2 FERM–kinase (2 μM). Pyk2 was incubated with ATP for 15 min at 37 °C to generate pY402 Pyk2. Autoinhibited WT Src was a product of incubation with 5 μM CSK and ATP for 10 min. YopH (0.3 μM) was added for 3.5 hr to dephosphorylate Y419 (**Fig. S4**). After 3.5 hr, 2 mM sodium orthovanadate was added to inhibit phosphatase activity. Open or autoinhibited WT Src was combined with pY402 Pyk2 (2 μM) and the reaction was monitored over a 60 min time course. Tyrosine phosphorylation of the Pyk2 activation loop was detected via Western blotting using Pyk2 pY579/pY580-specific primary antibody. (C) Activation loop phosphorylation of pY402 Pyk2 was quantified by densitometry from independent reactions (n = 3) at 60 min with either open or autoinhibited WT Src (**Fig. S4B**). (D) Kinase activity time course of autoinhibited WT Src (2 μM) with Pyk2 FERM–kinase (2 μM). Pyk2 was incubated for 15 min at 37 °C to generate pY402 Pyk2. WT pY530 Src was autoinhibited as described in B. Phosphorylation of the Src activation loop was detected via Western blotting using Src pY419-specific primary antibody. *Insets:* Keys to immunoblotting antibody recognition sites in A-C and D, respectively.

Next, we hypothesized that pY402 Pyk2 may promote efficient Src activation loop phosphorylation via scaffolding interactions that relieve the autoinhibitory Src conformation. Consequently, scaffolded Src could phosphorylate the Pyk2 activation loop (**Fig. 5B**), conferring Pyk2 the productive activation state for phosphorylation of the Src activation loop. To determine if Pyk2 also promotes the reciprocal Src activation loop phosphorylation, autoinhibited Src with or without pY402 Pyk2 was monitored for Src activation loop phosphorylation (**Fig. 5D**). Intriguingly, autoinhibited Src with or without pY402 Pyk2 exhibited similar rates of robust pY419 phosphorylation. As expected, no Src activation loop phosphorylation is detectable in a control repeated with Y419F Src (**Fig. S7**). Taken together, Src C-terminal phosphorylation sharply restricts kinase activity targeting the Pyk2 activation loop unless pY402 is available for scaffolding interaction. However, the C-terminal phosphorylation does not preclude autophosphorylation of the Src activation loop.

## Discussion

Src and Pyk2 undergo multistep conformational rearrangements that culminate in the activation of both kinases (**Fig. 6**). In this work, we reconstituted the Pyk2/Src system with defined regulatory states to dissect the mechanistic requirements for mutual activation loop phosphorylation. Additionally, we provide direct evidence of Pyk2-mediated conformational activation of Src. Taken together, several mechanistic themes emerge. First, activation loop phosphorylation and conformational competence for phosphoacceptor engagement are separable outputs. Disentangling the two clarifies the reciprocal regulation of the Pyk2–Src complex and reconciles observations that initially appear contradictory. Ultimately, our findings refine the mechanistic model for coordinated, reciprocal activation control between Src and Pyk2.

**Figure 6.**
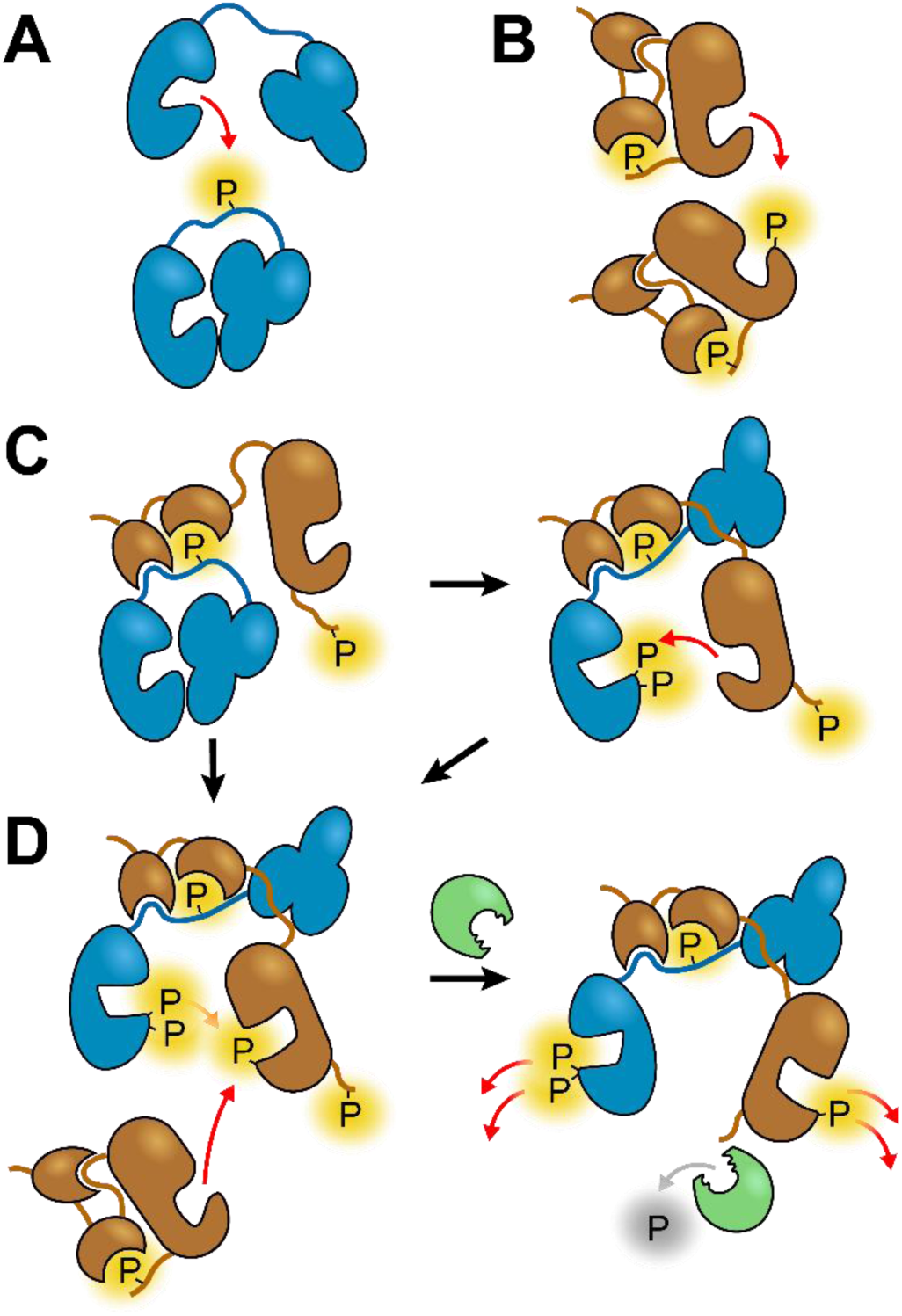
Proposed model for scaffolding-gated reciprocal regulation of Pyk2 and Src. Cartoon schematic of the ordered conformational and phosphorylation events that couple Pyk2 (blue) and Src (brown) activation. (A) Basal activity, autoinhibited Pyk2 is capable of autophosphorylation in trans to generate pY402 at the FERM—kinase linker. The proposed Ca^2+^-induced clustering that accelerates Pyk2 autophosphorylation is not shown. (B) Autoinhibited Src is capable of autophosphorylation in trans, but activation loop phosphorylation (pY419) alone does not confer full activity. Intramolecular SH3/SH2 engagement constrains access to other substrates. (C) Autophosphorylated Pyk2 presents tandem PRR and pY402 ligands that compete for intramolecular Src SH3/SH2 engagement, imparting conformational relief of Src autoinhibition independent of Src activation loop phosphorylation status. Unconstrained Src can phosphorylate the Pyk2 activation loop tyrosines (pY579/pY580) via an ordered, self-primed mechanism. (D) Active Pyk2 can reciprocally phosphorylate Src, but the activity is redundant with Src autophosphorylation. The open conformation of Pyk2-scaffolded Src exposes the C-terminal tail pY530 to co-localized phosphatases, committing both Src and Pyk2 to stable, maximal activity for downstream phosphorylation.

Src phosphorylates the Pyk2 activation loop with defined residue preference, favoring Y580 over Y579 (**Fig. 1**). As Pyk2 exhibits negligible activation loop autophosphorylation (**Fig. S1, S2**), Src is responsible for phosphorylation of both tandem activation loop tyrosines. Nevertheless, efficient phosphorylation of Y579 apparently requires prior phosphorylation of Y580. The Src preference for initial Y580 monophosphorylation is consistent with a sequence recognition register aligning a preferred bulky residue (Y) at position –1 and a preferred acidic residue (E) at position – 3 (62). We propose that Src engages in ordered, stepwise phosphorylation of the Pyk2 activation loop, in which monophosphorylation of Y580 reshapes local electrostatics and sterics to prime subsequent phosphorylation of Y579. Interestingly, this observation agrees well with recent work screening the sequence determinants of tandem phosphotyrosine specificity in Src and Abl kinases (63). Whereas Abl selectively engages phosphotyrosines neighboring N-terminal to the phosphoacceptor site (pYY), Src prefers phosphotyrosines immediately C-terminal to the target tyrosine (YpY). Within the Pyk2 activation loop, Src first generates pY580, which serves as a recognition mark to enable Src to complete phosphorylation of upstream residue Y579. As such, the Pyk2 activation loop serves as a self-priming Src substrate, where Src functions as both “reader” and “writer” of activation loop phosphotyrosines (63).

We also found that fully active (pY579/pY580) Pyk2 can directly phosphorylate the Src activation loop (Y419), but basal state Pyk2 cannot (**Fig. 2**). The reciprocal Pyk2/Src phosphorylation is efficient, albeit less rapid than Src autophosphorylation, and likely promoted by the close apposition of the two activation loops during complex formation. Notably, assembling the scaffolded complex with WT Src did not accentuate Src activation loop phosphorylation beyond the level achieved by Src autophosphorylation alone (**Fig. 5D**). This observation ostensibly conflicts with the observation that Pyk2 mediates robust Src activation loop phosphorylation (**Fig. 2**). We reconcile the two results by inferring that Src activation loop phosphorylation is dominated by rapid, autocatalytic trans autophosphorylation. The Src autoinhibited state remains relatively dynamic, sampling active-like conformations (64). A small Src subpopulation that manages autophosphorylation from the autoinhibited state quickly propagates activation loop phosphorylation independent of Pyk2. Therefore, while Pyk2 is competent for Src activation loop phosphorylation (**Fig. 2**), the activity is functionally redundant when high local concentrations of Src are available for autophosphorylation in trans. Rather, the mechanistic contribution of Pyk2 appears to be conformational control.

Autoinhibited (pY530) Src cannot phosphorylate a non-scaffolding substrate (e.g., Y402F Pyk2, **Fig. 4A and 5A**), yet autoinhibited Src does efficiently phosphorylate the Pyk2 activation loop when presented with the tandem SH3/SH2 ligands of pY402 Pyk2 (**Figs. 4B and 5B**). Because scaffolding rescues activity with non-phosphorylatable Y419F Src (**Fig. 4**), the phosphorylated Pyk2 linker relieves Src autoinhibition independent of Src activation loop phosphorylation. We interpret the Pyk2-mediated activation as scaffolding-based conformational activation of Src, in which the adjacent proline-rich region and pY402 outcompete intramolecular SH3/SH2 interactions that constrain Src in a catalytically incompetent state. The lag phase preceding Pyk2 activation loop phosphorylation by autoinhibited WT Src (**Fig. 5B**) suggests a gradual, scaffolding-dependent transition distinguishable from autocatalytic amplification.

The scaffolding-based conformational activation confers an important consequence for Src regulation. In the autoinhibited conformation, the C-terminal pY530 is protected from phosphatase activity (61), as we demonstrated by immunoblotting (**Fig. S5**). Relief of autoinhibition by Pyk2 scaffolding would expose pY530 to co-localized phosphatases, committing Src to a stable, high activity state. This model is reinforced by the allosteric coupling between the Src activation loop and the C-terminal phospho-motif. Because pY530 can restrain the kinase even when the activation loop is phosphorylated, activation loop status alone does not define the active state (60, 65). Src may thus carry pY419 while remaining conformationally autoinhibited, and only the combination of scaffold-mediated SH2/SH3 displacement and phosphatase-mediated pY530 removal yields maximal, committed activity for other substrate signaling partners. This distinction clarifies the apparent incongruity of robust pY530 Src autophosphorylation with the strict blockage of Pyk2 activation loop phosphorylation imposed by Src C-terminal phosphorylation.

The regulatory logic for Pyk2 is similarly layered. In the autoinhibited state, the Pyk2 FERM domain directly occludes the activation loop and substrate docking cleft while sequestering Y402 that serves as the Src-docking site (24, 25, 52). Relief of Pyk2 autoinhibition is upstream of Src engagement. To access the Pyk2 basal activity state, the FERM domain must transiently disengage to allow Pyk2 trans-autophosphorylation of Y402, a step that proceeds independent of Src (53). The resulting pY402 scaffolding site relieves Src autoinhibition, enabling Src to phosphorylate the Pyk2 activation loop, which Pyk2 cannot achieve alone (**Fig. S1**). Scaffolding may also allow Src to outcompete the tethered FERM domain for access to the Pyk2 activation loop. The fully phosphorylated Pyk2 activation loop is then committed to maximal activation, as phosphorylation prevents re-engagement of the FERM inhibitory interaction (24, 53). The trigger for FERM release remains unresolved, but proposed mechanisms include FERM-mediated dimerization, Ca²⁺/calmodulin-induced clustering, and PSD-95 scaffolding (16, 17, 24, 26).

Taken together, our results support a model in which reciprocal Pyk2–Src activation is organized as a coincidence detector. Neither kinase reaches full output in isolation: Pyk2 is gated by FERM autoinhibition and the dependence on Src for activation loop phosphorylation, while Src is gated by intramolecular SH2/SH3 autoinhibition and C-terminal pY530. Each kinase confers a regulatory input the other requires, restricting maximal signaling to the assembled complex. In this model, activated Pyk2 functions less as a direct upstream kinase for the Src activation loop than as a conformational activator and scaffolding platform. Pyk2 relieves Src autoinhibition, exposes the Src C-terminus for phosphatases, and positions maximally activated Src to phosphorylate downstream effectors. The findings refine the longstanding view of Pyk2 and FAK as dual signaling enzymes and scaffolds, clarifying the ordered, reciprocal regulatory events underlying activation.

## Experimental Procedures

### Protein purification

#### Pyk2 FERM***–***kinase purification

Plasmid pNAM004, encoding Pyk2 FERM–kinase (UniprotKB entry Q14289, residues 20–692), was co-expressed with pESU009, encoding the YopH tyrosine phosphatase domain (UniprotKB entry P08538, residues 177-468), in NiCo21(DE3) cells. Purification of Pyk2 and variants was performed as previously described (53).

#### Pyk2 kinase purification

Plasmid pSGC-PTK2BA encoding the Pyk2 kinase domain (UniprotKB entry Q14289, residues 414–692) was co-expressed with YopH (pESU009) in NiCo21(DE3) cells. After protein expression (16 hr, 18 °C) and cell harvesting by centrifugation, cells were lysed by sonication in 50 mM Tris pH 8.0, 25 mM NaCl, 20 mM imidazole, 5 mM EDTA, 5% glycerol, 1 mM PMSF, 2.5 mM β-mercaptoethanol (βME) with bacterial protease inhibitor cocktail II (Research Products International). Lysate was cleared of cell debris by ultracentrifugation at 80,000 ×g (Beckman MLA-50 rotor) for 15 min at 4°C. Pyk2 kinase was affinity purified on a Ni-NTA affinity column (HisTrap HP 5 mL, Cytiva). The H_6_-TEV tag was cleaved via GFP-TEV-His_6_ protease treatment, and protease and cleaved tag was removed by passage through a subtractive Ni-NTA column. Enriched Pyk2 kinase was concentrated using a 10k MWCO PES Spin-X centrifugal concentrator (Corning). The concentrated sample was resolved through a gel filtration column (Superdex 200 10/300, GE Healthcare) pre-equilibrated in 50 mM HEPES pH 7.2, 150 mM NaCl, 5% glycerol, and 5 mM TCEP. Purified Pyk2 kinase domain was aliquoted, snap frozen in liquid N_2_, and stored at –80°C.

#### Src purification

Src (UniprotKB entry P12931, residues 1–536) encoded by pJC101 was co-expressed with pESU009 (YopH) in NiCo21(DE3) cells. Purification was performed as previously described (53).

#### FLAG-CSK purification

Plasmid pAAW034 encoding H_6_-SUMO-FLAG-CSK (UniprotKB entry P41240, residues 1–450) was expressed in NiCo21(DE3) cells for 16 hr at 18 °C. Cells were harvested by centrifugation and stored at –80°C until purification. Cells were thawed on ice and lysed by sonication in 50 mM Tris pH 8.0, 25 mM NaCl, 20 mM imidazole, 5 mM EDTA, 5% glycerol, 1 mM PMSF, 2.5 mM βME, supplemented with protease inhibitor cocktail II (Research Products International). Lysate was cleared of cell debris by ultracentrifugation at 80,000 ×g (Beckman MLA-50 rotor) for 20 min at 4°C. H_6_-SUMO-FLAG-CSK was captured on Ni-NTA (5 mL HisTrap, Cytiva) equilibrated with 50 mM Tris pH 8.0, 250 mM NaCl, 20 mM imidazole, 5% glycerol. His-tagged protein was eluted using 50 mM Tris pH 8.0, 150 mM NaCl, 300 mM imidazole, 5% glycerol. The H_6_-SUMO tag was cleaved through incubation with H_6_-Ulp1 protease. Sample was dialyzed into 50 mM Tris pH 8.0, 150 mM NaCl, 5% glycerol, 2.5 mM βME. Protease and H_6_-SUMO tag were removed by a subtractive passage through Ni-NTA resin. FLAG-CSK was concentrated using a 10k MWCO PES Spin-X centrifugal concentrator (Corning) and applied to a gel filtration column (Superdex 200 10/300, GE Healthcare) pre-equilibrated in 50 mM HEPES pH 7.2, 150 mM NaCl, 5% glycerol, and 5 mM TCEP. Purified proteins were aliquoted, snap frozen in liquid N_2_, and stored at –80°C.

#### Avi-YopH purification

Plasmid pEUS007, encoding Avi-tagged YopH (UniprotKB entry P08538, residues 177–468) was transformed and expressed in BL21(DE3) cells. Cells were harvested by centrifugation and stored at –80°C until purification. Cells were thawed on ice and lysed by sonication in 50 mM Tris pH 8.0, 25 mM NaCl, 25 mM imidazole, 5 mM EDTA, 5% glycerol, 1 mM PMSF, 2.5 mM βME. Lysate was cleared of cell debris by centrifugation at 20,000 ×g (Fiberlite F15-8×50c) for 40 min at 4°C. Avi-YopH purified using cation exchange chromatography using a HiTrap SP FF (5 mL, Cytiva) equilibrated with low salt buffer (25 mM HEPES pH 7.0, 20 mM NaCl, 5% glycerol, 2.5 mM βME). Proteins were resolved with a linear gradient from low to high salt buffer (50 mM HEPES pH 7.0, 500 mM NaCl, 5% glycerol). Enriched proteins were aliquoted, snap frozen in liquid N_2_, and stored at –80°C.

### Kinase Assays

Kinase reactions were performed in a Kinase Buffer composed of 50 mM HEPES pH 7.4, 150 mM NaCl, 8 mM MgCl_2_, 5% glycerol. Reactions were initiated by addition of ATP to a final concentration of 4 mM and quenched with 8.3 mM EDTA and 1.5% SDS (final concentrations). SDS was omitted from the quench for dot blotting assays. Phosphotyrosine levels were measured via Western or dot blotting on 0.2 µm nitrocellulose (Cytiva Amersham). Briefly, blots were blocked with BSA and probed with site-specific primary antibodies (Table 1) followed by goat anti-rabbit secondary antibody conjugated to horse radish peroxidase (Invitrogen, A16096). Blots were imaged with enhanced chemiluminescent substrate (Pierce).

**Table 1.** Antibodies for site-specific anti-phosphotyrosine Western and dot blot analysis.

| IB Label | Name | Vendor (Product Number) |
| --- | --- | --- |
| Pyk2 pY579/pY580 | Phospho-PYK2 (Tyr579, Tyr580) polyclonal | Invitrogen (44-636G) |
| Pyk2 pY402 | Anti-phospho-PTK2B-pY402 polyclonal | Sigma-Aldrich (SAB4300173) |
| Src pY419 | Phospho-SRC (Tyr416) polyclonal<sup>1</sup> | Invitrogen (PA5-97364) |
| Src pY419 monoclonal | Phospho-Src Family (Tyr416) monoclonal (D49G4)<sup>1</sup> | Cell Signaling Technologies (6943) |
| Src pY530 | Phospho-SRC (Tyr527) polyclonal<sup>1</sup> | Invitrogen (PA5-117390) |

<sup>1</sup>Site-specific commercial antibodies for Src use avian residue numbering, whereas Src phospho-site residues reported throughout this work are labeled with human residue numbering.

#### Src phosphorylation of Pyk2 activation loop variants

Pyk2 FERM–kinase variants (WT, Y579T, Y580T, or Y579T/Y580T) were incubated in Kinase Buffer with ATP at 37°C for 120 min to fully phosphorylate Y402. Pyk2 variants (0.5 μM) and Src (0.2 μM) were incubated in Kinase Buffer with 4 mM ATP and sampled throughout a 60 min time course. Triplicate reaction time points were measured at 5 min from three independent reactions. Phosphorylation of the Pyk2 activation loop was assessed via Western blotting using anti-Pyk2 pY579/pY580 primary antibody.

#### Src Y419F phosphorylation of Pyk2

Pyk2 FERM–kinase was incubated in Kinase Buffer with 4 mM ATP for 15 min at 37°C to generate pY402 Pyk2 (0.5 μM). Either Src WT (0.5 μM) or Src Y419F (0.5 μM) were added to the pY402 Pyk2 pre-incubation, and the Src reaction was monitored throughout a 60 min time course. Phosphorylation of the Pyk2 activation loop was assessed via Western blotting using anti-Pyk2 pY579/pY580.

Kinase dead K457A Pyk2 FERM–kinase (0.5 μM) was incubated in Kinase Buffer with 4 mM ATP and either WT Src (0.5 μM) or Y419F Src (0.5 μM). The kinase reaction was sampled over a 60 min time course. Phosphorylation of the Pyk2 autophosphorylation site was assessed via Western blotting using anti-Pyk2 pY402 primary antibody.

#### Pyk2 phosphorylation of Src

*Basal Pyk2* – Pyk2 FERM–kinase (1 μM) plus K298M Src (1 μM) or WT Src (1 μM) were incubated with 4 mM ATP in Kinase Buffer and monitored over a 60 min time course. Phosphorylation of the Src activation loop was assessed via Western blotting using anti-Src pY419 polyclonal primary antibody.

*Active Pyk2* – Fully phosphorylated pY402/pY579/pY580 Pyk2 FERM–kinase was obtained via pre-incubation with Src. Pyk2 (23 μM) was incubated with Src (0.2 μM) and 4 mM ATP in Kinase Buffer for 180 min. To control for the background of residual Src used to activate pY579/pY580 Pyk2 FERM–kinase, a mock reaction was performed omitting Pyk2 FERM–kinase with the same overall dilution of catalytic Src (0.2 µM initial, 8 nM final). The fully phosphorylated Pyk2 stock was added to a final concentration of 1 μM to kinase dead K298M or WT Src (1 μM) and 4 mM ATP, and phosphotransfer was monitored at various time points over 60 min. For triplicate data points, three independent reaction replicates were assessed at 15 min. Phosphorylation of the Src activation loop was assessed via Western blotting using anti-Src pY419 polyclonal antibody.

#### Src activity in open vs. autoinhibited conformations

To phosphorylate Y530 of Src, FLAG-CSK (5 µM) was incubated with either WT or Y419F Src plus 4 mM ATP for 10 min in 50 mM HEPES pH 7.4, 50 mM NaCl, 8 mM MgCl_2_, 5% glycerol. Avi-YopH (0.3 μM) was added to the reaction and incubated for 3.5 hours to dephosphorylate the Src activation loop (pY419) and subsequently inhibited with the addition of 2 mM sodium orthovanadate. WT or Y419F Src WT (2 μM) in either open or autoinhibited (pY530) conformation were combined with pY402 Pyk2 (2 μM), and the reaction was monitored for 60 min. Phosphorylation of the Pyk2 activation loop was assessed via Western blotting using anti-Pyk2 pY579/pY580 primary antibody. Src phosphorylation was assessed via Western blotting using anti-Src pY419 monoclonal primary antibody and anti-Src pY530 primary antibody.

#### Active Pyk2 phosphorylation of Pyk2 activation loop

Fully phosphorylated pY402/pY579/pY580 Pyk2 FERM–kinase was generated via incubation with Src, as described above. The fully phosphorylated Pyk2 stock was added to a final concentration of 1 μM to Pyk2 kinase domain (1 μM), and phosphotransfer was initiated by addition of ATP (4 mM). A control reaction for background Src was performed in parallel, omitting Pyk2 FERM–kinase, with the same overall dilution of catalytic Src (0.2 µM initial, 8 nM final) added to Pyk2 kinase domain (1 µM). The reaction was sampled over a 60 min time course. Phosphorylation of the Pyk2 kinase domain activation loop was assessed via Western blotting using anti-Pyk2 pY579/pY580.

### Intact Protein Mass Spectrometry

Pyk2 FERM−kinase (1.5 µM) was pre-incubated with 4 mM ATP and 2 mM orthovanadate for 2 hr (37 °C) to generate pY402 Pyk2. Pyk2 activation loop phosphorylation was assessed after addition of Src (0.5 µM) and 5 min incubation at 37°C. Reactions were quenched by addition of EDTA to 8.3 mM final concentration. Quenched samples were injected (2 μL) into an ACQUITY UPLC H-class instrument coupled in line to an ESI-Q-TOF Synapt G2-Si mass spectrometer (Waters). Mobile phases consisted of solvents A (HPLC-grade aqueous 0.1% formic acid) and B (HPLC-grade acetonitrile and 0.1% formic acid). Exchange samples were desalted on a C4 column (5 μm, 1 mm × 50 mm; Restek) with 5% solvent B for 1 min at a flow rate of 400 μL/ min. Intact protein was eluted with a gradient ramp from 5% to 100% solvent B over 8 min. Mass spectra were collected in positive ion, MS continuum, resolution mode with an m/z range of 300−4000. The intact mass of each protein species was deconvoluted from multiple charge states using the MaxEnt1 function of MassLynx (Waters).

## Acknowledgements

We thank Nicola Burgess-Brown for the gift of pSGC-PTK2BA (Addgene plasmid #42401). We thank Joel Nott of the Iowa State University Protein Facility for mass spectrometry support.

## Author Contributions

Conceptualization, A.W., T.M.P.Z., and E.S.U.; methodology, A.W. and T.M.P.Z.; investigation, A.W., K.M.Z., T.M.P.Z., and K.G.R.B.; writing—original draft, A.W.; writing—review & editing, E.S.U and A.W.; funding acquisition, E.S.U.

## Abbreviations

Pyk2: Proline-rich tyrosine kinase 2
FAK: focal adhesion kinase
FERM: protein 4.1/ezrin/radixin/moesin
FAT: focal adhesion targeting
PRR: Proline-rich region
IB: immunoblot
SFK: Src family kinase
CSK: C-terminal Src kinase
FK: FERM–kinase

## Supporting information

**Figure S1.**
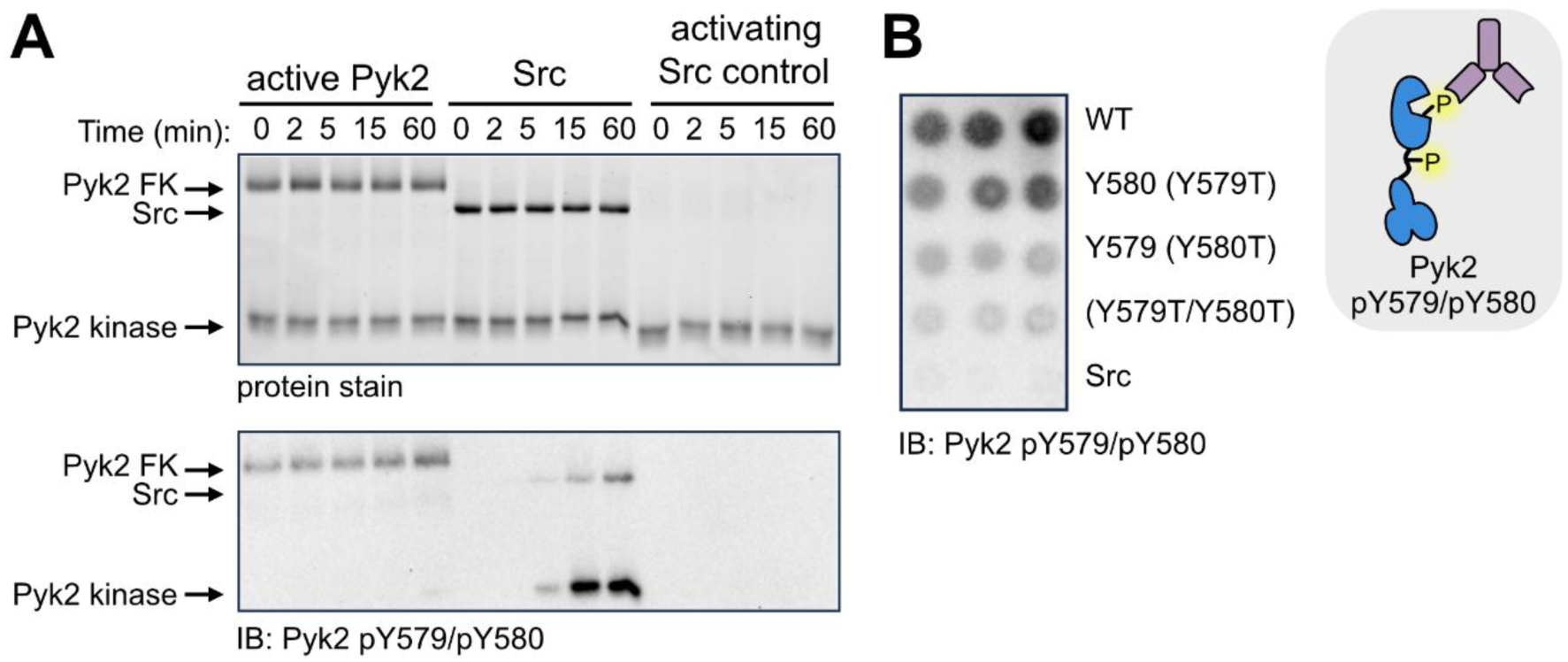
Kinase specificity in Pyk2 activation loop phosphorylation. (A) Activated Pyk2 (pY579/pY580) does not function in trans to phosphorylate other Pyk2 activation loops. Unphosphorylated Pyk2 kinase domain (residues 414–692) was incubated with either pY579/pY580 Pyk2 FERM–kinase (residues 20–692) or Src. Fully phosphorylated pY579/pY580 Pyk2 FERM–kinase (23 μM) was generated via pre-incubation with Src (0.2 μM) and ATP for 180 min. Activation loop phosphorylation of initially unphosphorylated Pyk2 kinase domain (1 µM) was monitored over a 60 min time course with pY579/pY580 Pyk2 FERM–kinase (1 µM, left) or Src (1 µM, center). To control for the background of residual Src used to activate pY579/pY580 Pyk2 FERM–kinase, a mock reaction was performed omitting Pyk2 FERM–kinase with the same overall dilution of catalytic Src (right). Tyrosine phosphorylation of the Pyk2 activation loop was detected via Western blotting using Pyk2 pY579/pY580-specific primary antibody. *Inset:* Key to immunoblotting antibody recognition sites in A and B. (B) Activation loop phosphorylation of Pyk2 FERM–kinase variants (0.5 μM) in triplicate at 5 min in the presence of Src (0.2 μM). Tyrosine phosphorylation of the Pyk2 activation loop was detected via dot blotting using Pyk2 pY579/pY580-specific primary antibody.

**Figure S2.**
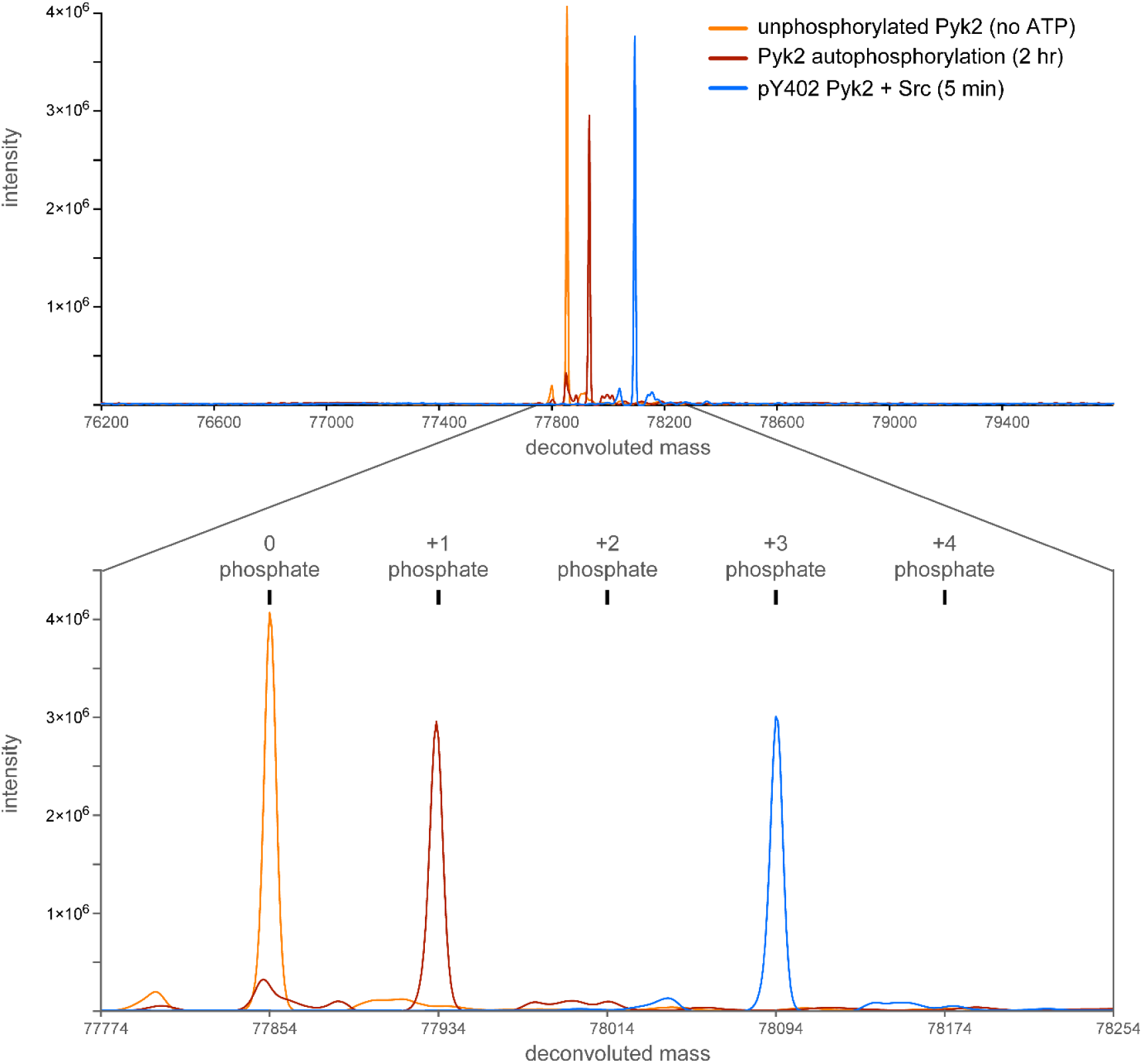
Intact protein mass spectrometry supports dual phosphorylation of the Pyk2 activation loop by Src. Deconvoluted intact mass spectra of Pyk2 FERM–kinase were collected at successive stages of phosphorylation. Prior to ATP incubation (orange), Pyk2 (1.5 µM) exhibits a single species consistent with the unphosphorylated protein (expected mass: 77,854 Da). Following a pre-incubation with 4 mM ATP (37 °C, 2 hrs), the Pyk2 spectrum (brown) exhibits a mass shift (ΔM = + 80 Da) consistent with addition of a single phosphate (expected monophosphorylated Pyk2 mass: 77,934 Da). Addition of Src (0.5 µM) converted the monophosphorylated Pyk2 spectrum (blue) to a new mass consistent with two additional phosphorylations (ΔM = + 240 Da relative to unphosphorylated, expected triphosphorylated Pyk2 mass: 78,094 Da) within 5 min. Expected masses are reported as average mass values of Pyk2 residues 20–692.

**Figure S3.**
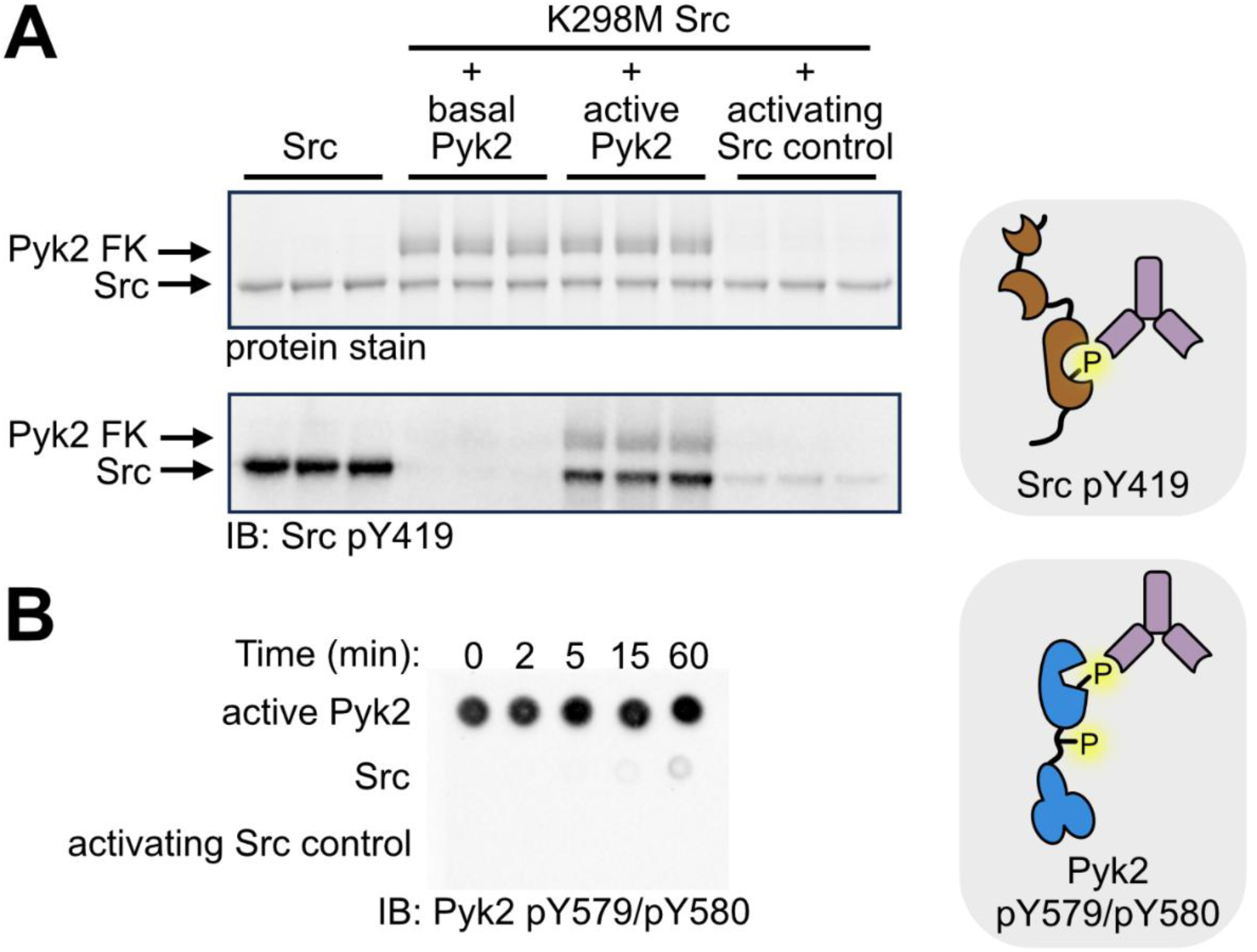
Active Pyk2 phosphorylates Src Y419. (A) Comparison of Src activation loop phosphorylation mediated by basal (pY402) Pyk2, active (pY579/pY580) Pyk2, or Src autophosphorylation. Basal Pyk2 or active Pyk2 (1 μM) were incubated with K298M Src (1 μM). WT Src autophosphorylation (1 µM) was assessed without Pyk2 (leftmost triplicates). To control for the background of residual Src used to activate pY579/pY580 Pyk2, a mock reaction was performed omitting Pyk2 with the same overall dilution of catalytic Src (rightmost triplicate). Triplicate, independent reactions were quenched and analyzed at 15 min. Phosphorylation was detected via Western blotting with Src pY419-specific primary antibody. (B) Dot blot assessing activation loop phosphorylation state of Pyk2 throughout the reaction time course as detected by Pyk2 pY579/pY580-specific primary antibody. Src and diluted catalytic Src control reactions are included to confirm antibody specificity. *Insets:* Keys to immunoblotting antibody recognition sites in A and B, respectively.

**Figure S4.**
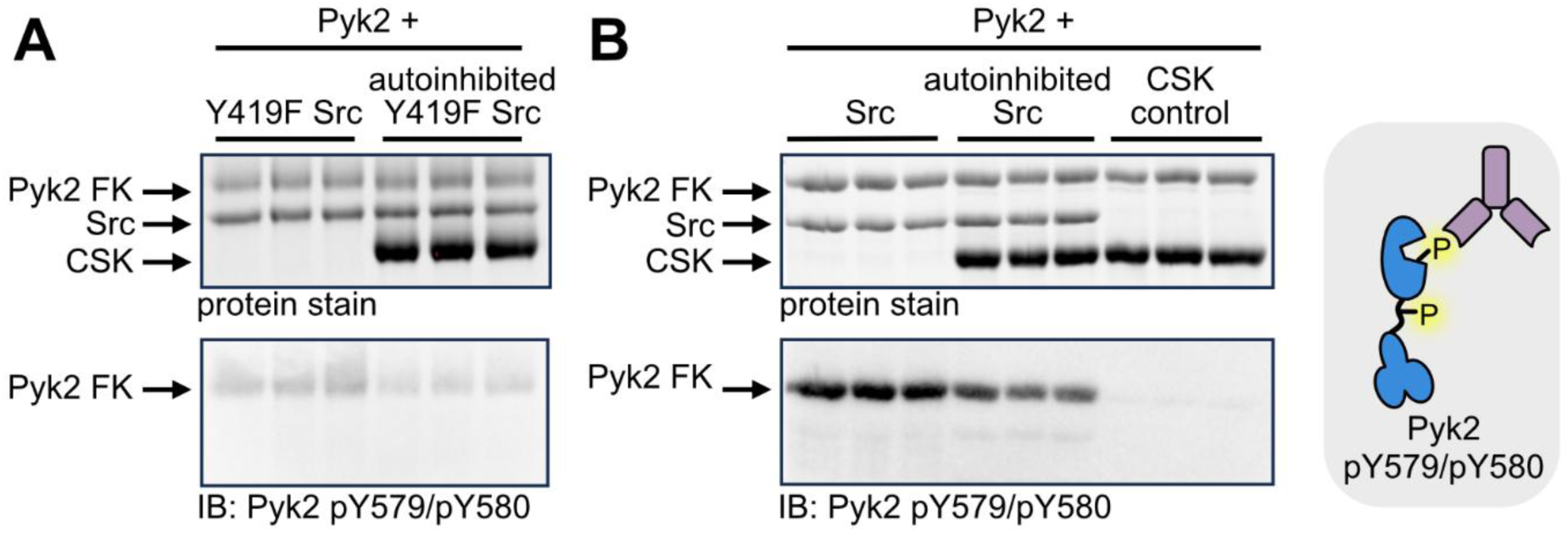
Pyk2 pY402 scaffolding outcompetes Src autoinhibition. (A) Kinase activity of open (unphosphorylated) or autoinhibited (pY530) Y419F Src (2 μM) with pY402 Pyk2 FERM–kinase (2 μM). Autoinhibited Y419F Src (2 μM) was generated by incubation with 5 μM CSK and ATP for 10 min. Basal or autoinhibited Y419F Src was combined with pY402 Pyk2 (2 μM), and kinase activity was assessed at 60 min for three independent reaction replicates. Phosphorylation of the Pyk2 activation loop was detected via Western blotting using Pyk2 pY579/pY580-specific primary antibody. (B) Kinase activity time course of open or autoinhibited WT Src (2 μM) with Pyk2 FERM–kinase (2 μM). Pyk2 was incubated with ATP for 15 min at 37°C to generate pY402 Pyk2. Autoinhibited WT Src was a product of incubation with 5 μM CSK and ATP for 10 min. YopH (0.3 μM) was added for 3.5 hr to dephosphorylate Y419. After 3.5 hr, 2 mM sodium orthovanadate was added to inhibit phosphatase activity. Open or autoinhibited Src WT was combined with pY402 Pyk2 (2 μM) and the reaction was assessed at 60 min for three independent reaction replicates. Tyrosine phosphorylation of the Pyk2 activation loop was detected via Western blotting using Pyk2 pY579/pY580-specific primary antibody. *Inset:* Key to immunoblotting antibody recognition sites in A and B.

**Figure S5.**
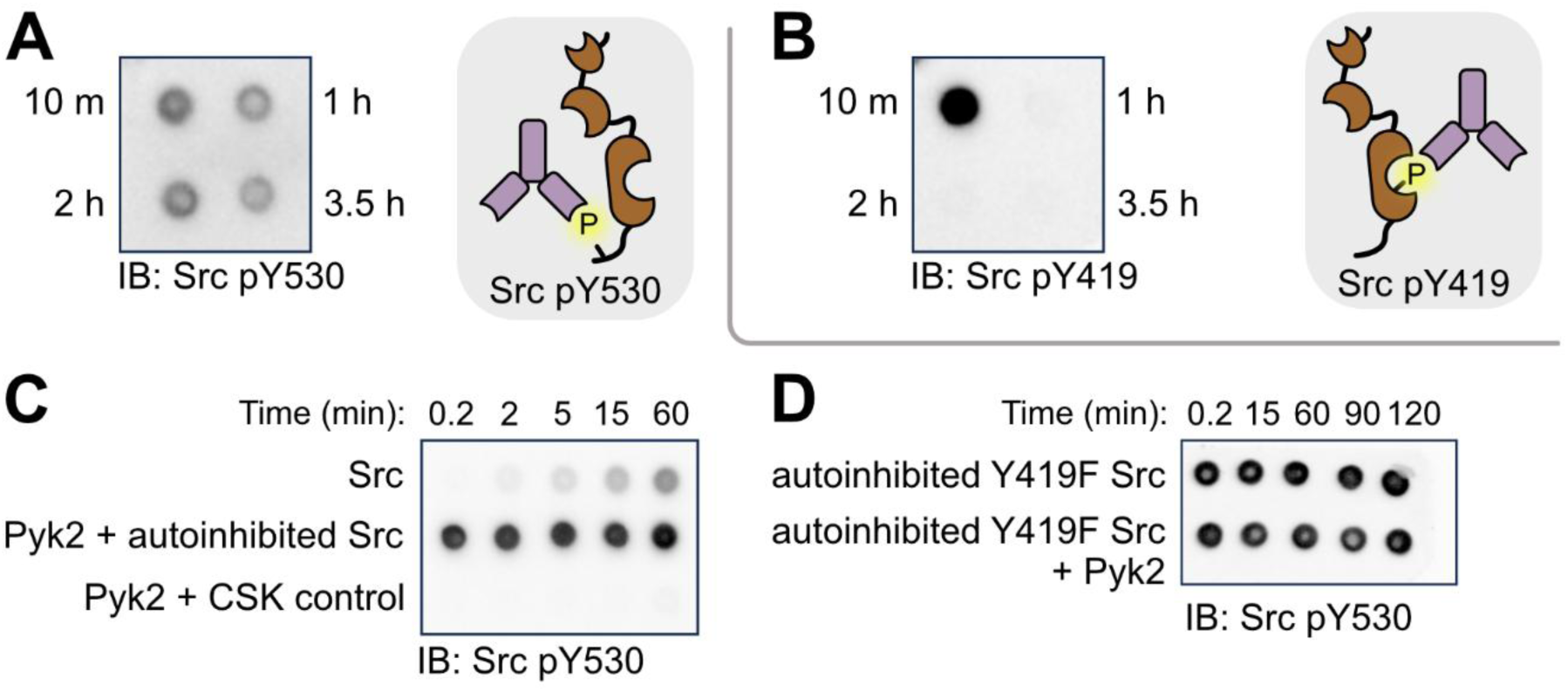
Src inhibition by CSK. (A, B) Parallel dot blots monitoring Src Y530 phosphorylation (A) and Src Y419 phosphorylation (B) after CSK incubation (10 min) followed by incubation with YopH to remove pY419 (1 h, 2 h, 3.5 h). Tyrosine phosphorylation of the Src C-terminus (A) was detected via dot blotting using Src pY530-specific primary antibody. Tyrosine phosphorylation of the Src activation loop (B) was detected via dot blotting using Src pY419-specific monoclonal primary antibody. (C) Time courses monitoring Src Y530 phosphorylation state of open (unphosphorylated) Src, Pyk2 and autoinhibited (pY530) Src, or Pyk2 and CSK. Each reaction includes 4 mM ATP. Tyrosine phosphorylation of the Src C-terminus was detected via dot blotting using Src pY530-specific primary antibody. (D) Time courses monitoring Src Y530 phosphorylation state for autoinhibited (pY530) Src Y419F with and without Pyk2. Tyrosine phosphorylation of the Src C-terminus was detected via dot blotting using Src pY530-specific primary antibody. *Inset:* Keys to immunoblotting antibody recognition sites in A, C, and D (Src pY530) and B (Src pY419), respectively.

**Figure S6.**
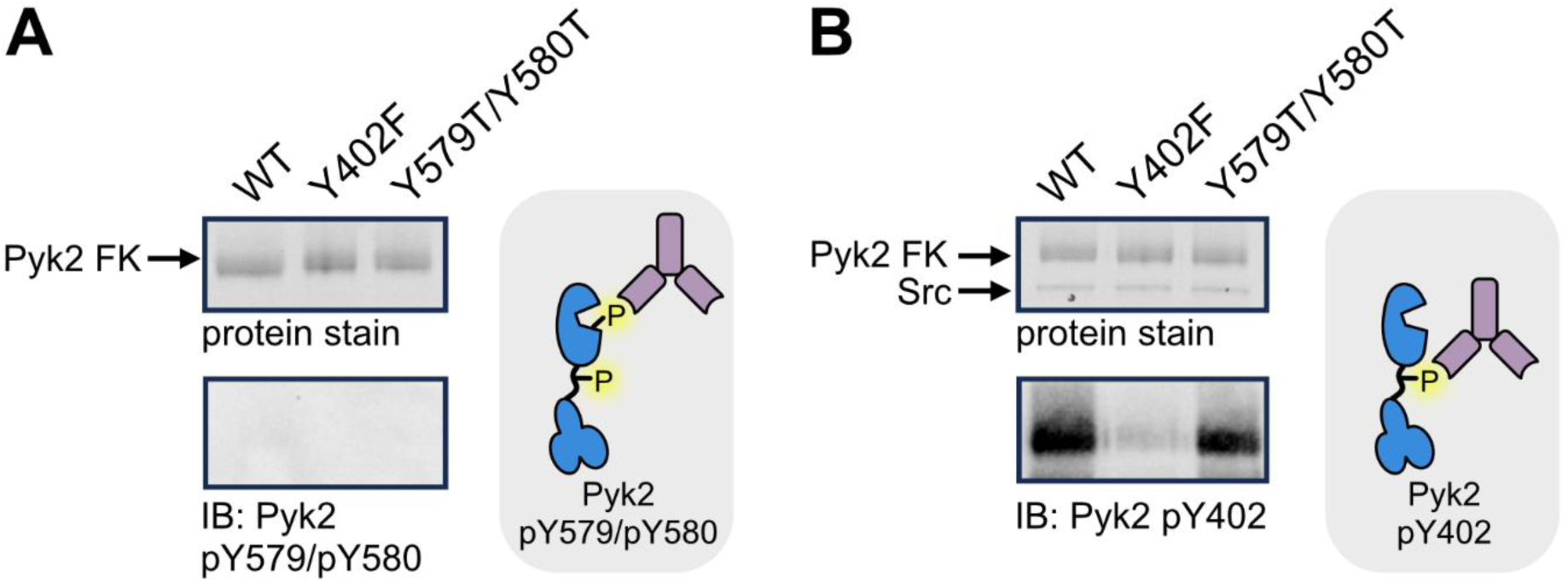
Testing Pyk2 and Src phosphosite targeting specificity. (A) Activation loop autophosphorylation of WT, Y402F, and Y579T/Y580T Pyk2 FERM–kinase (1 μM). Kinase variants were incubated with 4 mM ATP for 60 min at 37 °C. Phosphorylation of the Pyk2 activation loop was probed via Western blotting using Pyk2 pY579/pY580-specific primary antibody. (B) Phosphorylation of the FERM—kinase linker of WT, Y402F, and Y579T/Y580T Pyk2 FERM-kinase (1 μM) with basal state WT Src (0.2 μM) at 60 min. Tyrosine phosphorylation of the Pyk2 activation loop was probed via Western blotting using Pyk2 pY402-specific primary antibody. *Insets:* Keys to immunoblotting antibody recognition sites in A and B, respectively.

**Figure S7.**
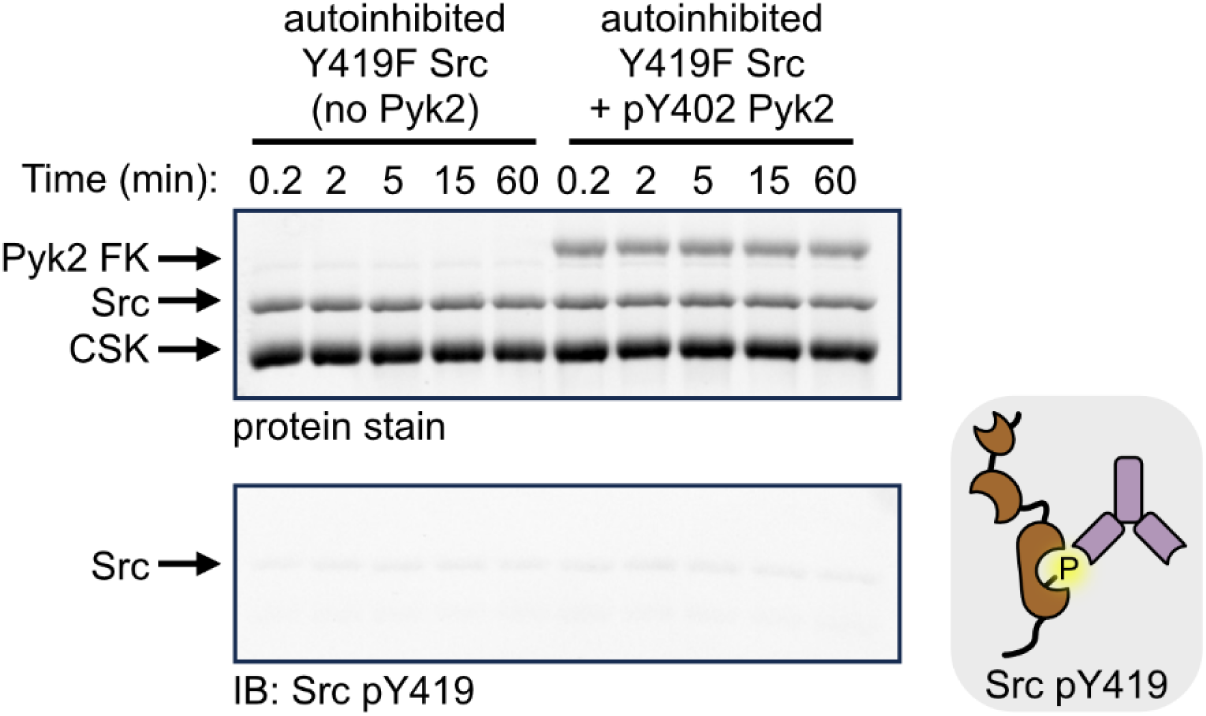
Testing for autoinhibited Src phosphotransfer background. To test for detectable phosphotransfer background of autoinhibited Src, pY530/Y419F Src (2 μM) was incubated in the presence or absence of pY402 Pyk2 FERM–kinase (2 μM) with 4 mM ATP. Autoinhibited, pY530/Y419F Src was generated as described in **Fig. 4**. Phosphorylation of the Src activation loop was probed via Western blotting using Src pY419-specific monoclonal primary antibody. *Inset:* Key to immunoblotting antibody recognition site.

