## Supporting Information Figures S1-S7 for "Deciphering the reciprocal regulation of the Pyk2–Src activation complex"

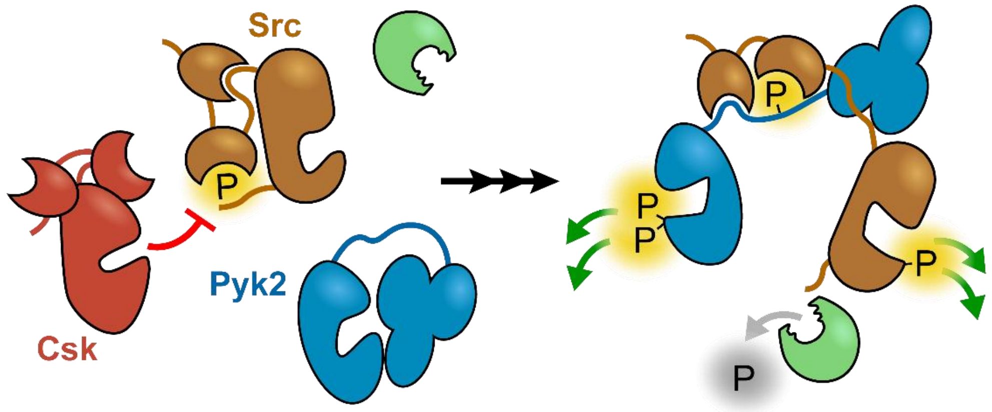

**Supporting information**

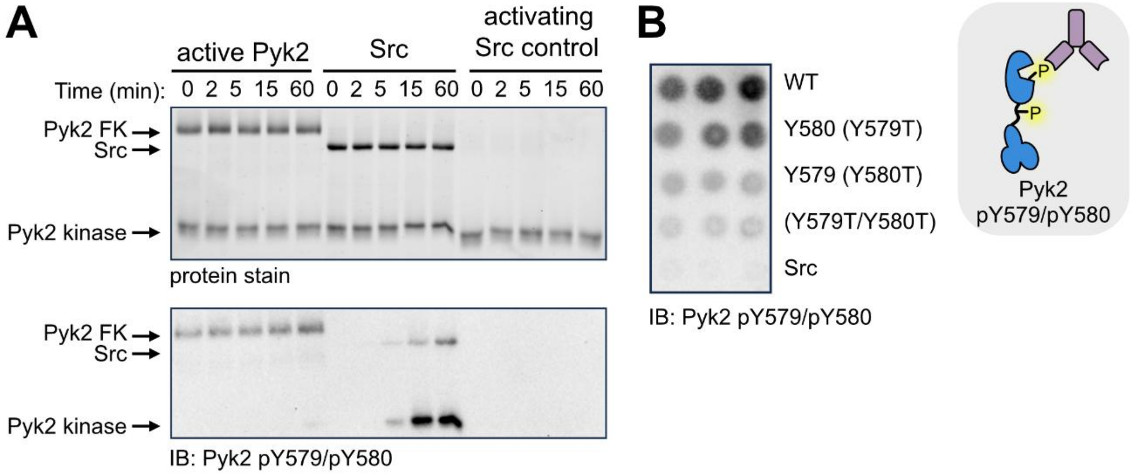


**Figure S1. Kinase specificity in Pyk2 activation loop phosphorylation**
(A) Activated Pyk2 (pY579/pY580) does not function in trans to phosphorylate other Pyk2 activation loops. Unphosphorylated Pyk2 kinase domain (residues 414–692) was incubated with either pY579/pY580 Pyk2 FERM–kinase (residues 20–692) or Src. Fully phosphorylated pY579/pY580 Pyk2 FERM–kinase (23 μM) was generated via pre-incubation with Src (0.2 μM) and ATP for 180 min. Activation loop phosphorylation of initially unphosphorylated Pyk2 kinase domain (1 μM) was monitored over a 60 min time course with pY579/pY580 Pyk2 FERM–kinase (1 μM, left) or Src (1 μM, center). To control for the background of residual Src used to activate pY579/pY580 Pyk2 FERM–kinase, a mock reaction was performed omitting Pyk2 FERM–kinase with the same overall dilution of catalytic Src (right). Tyrosine phosphorylation of the Pyk2 activation loop was detected via Western blotting using Pyk2 pY579/pY580-specific primary antibody. *Inset:* Key to immunoblotting antibody recognition sites in A and B. (B) Activation loop phosphorylation of Pyk2 FERM–kinase variants (0.5 μM) in triplicate at 5 min in the presence of Src (0.2 μM). Tyrosine phosphorylation of the Pyk2 activation loop was detected via dot blotting using Pyk2 pY579/pY580-specific primary antibody.

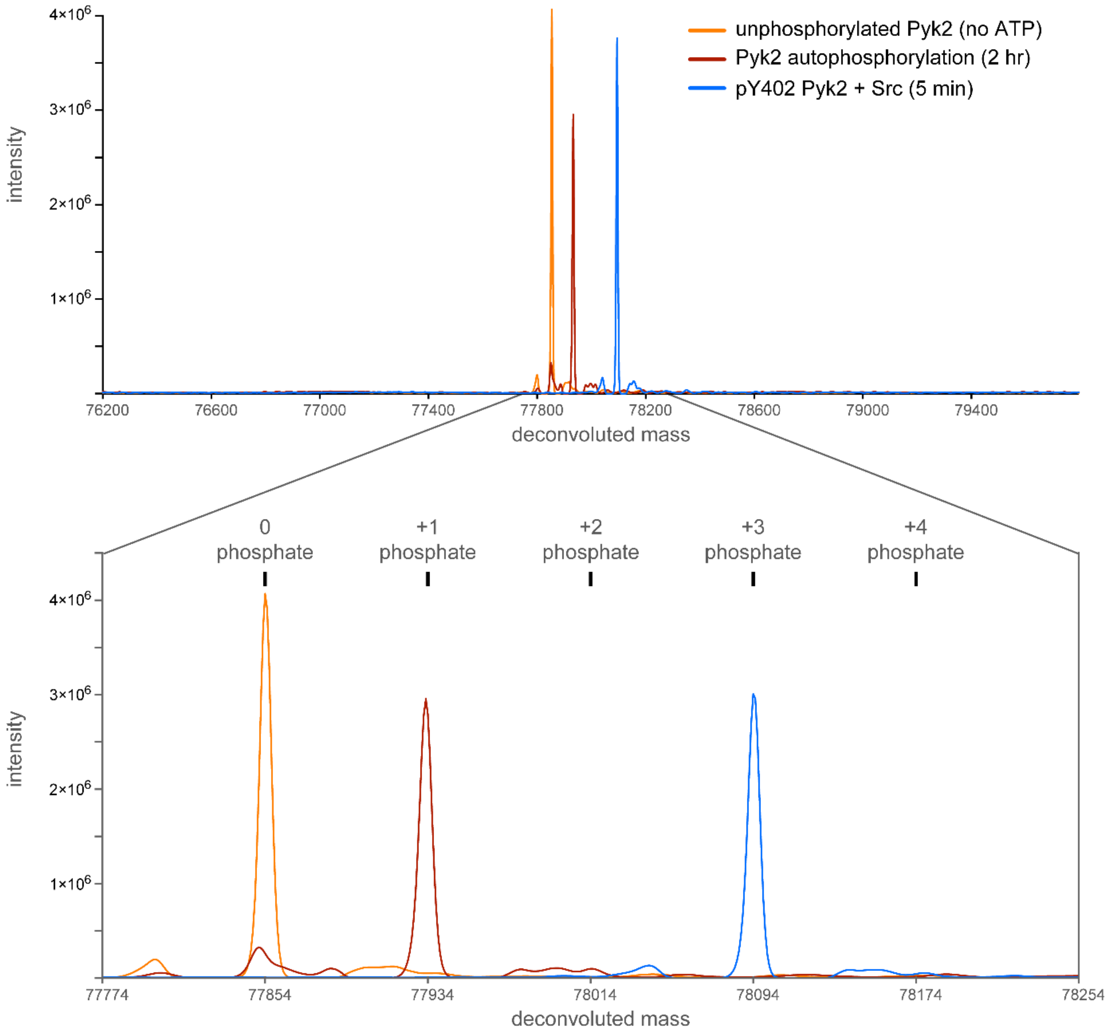


**Figure S2.** Intact protein mass spectrometry supports dual phosphorylation of the Pyk2 activation loop by Src. Deconvoluted intact mass spectra of Pyk2 FERM–kinase were collected at successive stages of phosphorylation. Prior to ATP incubation (orange), Pyk2 (1.5 μM) exhibits a single species consistent with the unphosphorylated protein (expected mass: 77,854 Da). Following a pre-incubation with 4 mM ATP (37 °C, 2 hrs), the Pyk2 spectrum (brown) exhibits a mass shift (ΔM = + 80 Da) consistent with addition of a single phosphate (expected monophosphorylated Pyk2 mass: 77,934 Da). Addition of Src (0.5 μM) converted the monophosphorylated Pyk2 spectrum (blue) to a new mass consistent with two additional phosphorylations (ΔM = + 240 Da relative to unphosphorylated, expected triphosphorylated Pyk2 mass: 78,094 Da) within 5 min. Expected masses are reported as average mass values of Pyk2 residues 20–692.

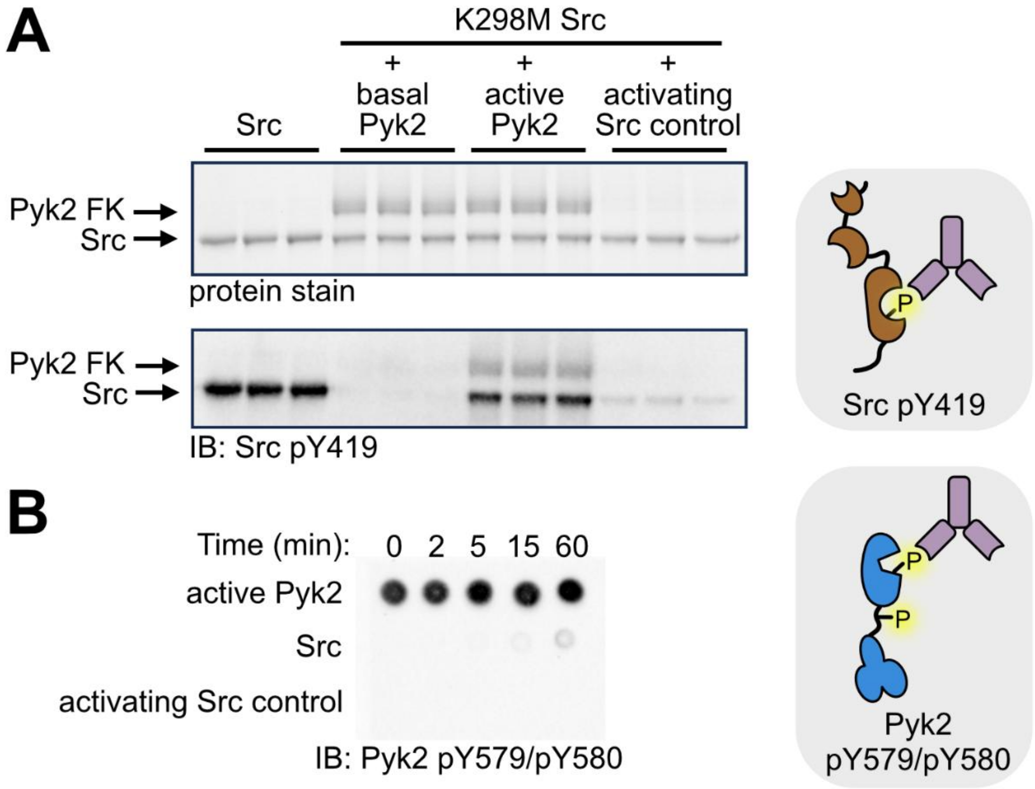


**Figure S3. Active Pyk2 phosphorylates Src Y419**

(A) Comparison of Src activation loop phosphorylation mediated by basal (pY402) Pyk2, active (pY579/pY580) Pyk2, or Src autophosphorylation. Basal Pyk2 or active Pyk2 (1 µM) were incubated with K298M Src (1 µM). WT Src autophosphorylation (1 µM) was assessed without Pyk2 (leftmost triplicates). To control for the background of residual Src used to activate pY579/pY580 Pyk2, a mock reaction was performed omitting Pyk2 with the same overall dilution of catalytic Src (rightmost triplicate). Triplicate, independent reactions were quenched and analyzed at 15 min. Phosphorylation was detected via Western blotting with Src pY419-specific primary antibody. (B) Dot blot assessing activation loop phosphorylation state of Pyk2 throughout the reaction time course as detected by Pyk2 pY579/pY580-specific primary antibody. Src and diluted catalytic Src control reactions are included to confirm antibody specificity. *Insets:* Keys to immunoblotting antibody recognition sites in A and B, respectively.

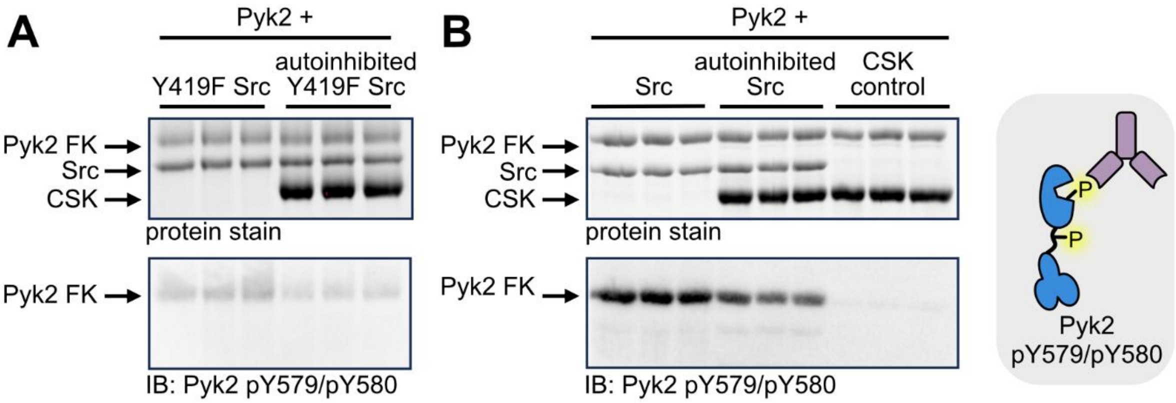


**Figure S4. Pyk2 pY402 scaffolding outcompetes Src autoinhibition**
(A) Kinase activity of open (unphosphorylated) or autoinhibited (pY530) Y419F Src (2 μM) with pY402 Pyk2 FERM–kinase (2 μM). Autoinhibited Y419F Src (2 μM) was generated by incubation with 5 μM CSK and ATP for 10 min. Basal or autoinhibited Y419F Src was combined with pY402 Pyk2 (2 μM), and kinase activity was assessed at 60 min for three independent reaction replicates. Phosphorylation of the Pyk2 activation loop was detected via Western blotting using Pyk2 pY579/pY580-specific primary antibody. (B) Kinase activity time course of open or autoinhibited WT Src (2 μM) with Pyk2 FERM–kinase (2 μM). Pyk2 was incubated with ATP for 15 min at 37 ˚C to generate pY402 Pyk2. Autoinhibited WT Src was a product of incubation with 5 μM CSK and ATP for 10 min. YopH (0.3 μM) was added for 3.5 hr to dephosphorylate Y419. After 3.5 hr, 2 mM sodium orthovanadate was added to inhibit phosphatase activity. Open or autoinhibited Src WT was combined with pY402 Pyk2 (2 μM) and the reaction was assessed at 60 min for three independent reaction replicates. Tyrosine phosphorylation of the Pyk2 activation loop was detected via Western blotting using Pyk2 pY579/pY580-specific primary antibody. *Inset:* Key to immunoblotting antibody recognition sites in A and B.

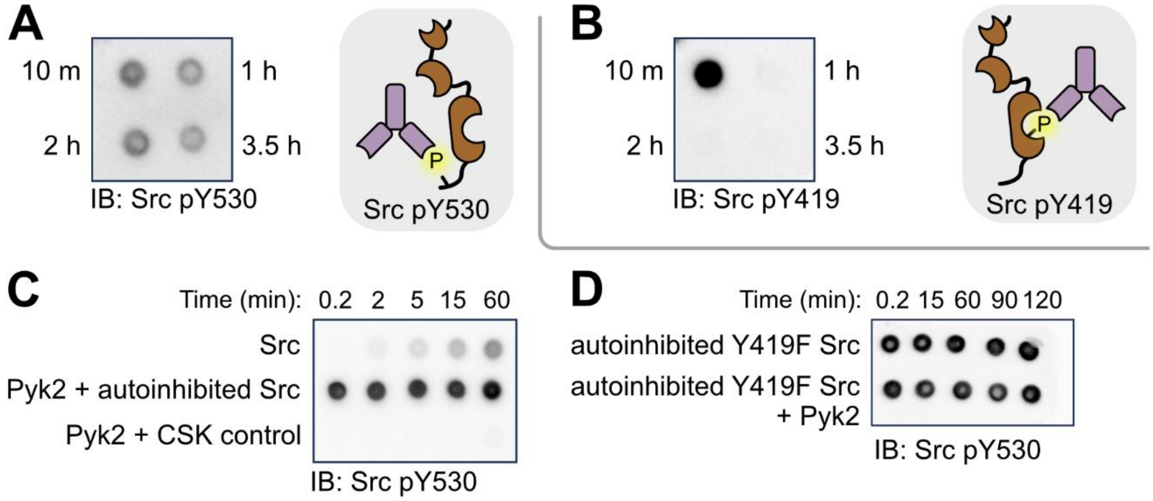


**Figure S5. Src inhibition by CSK**

(A, B) Parallel dot blots monitoring Src Y530 phosphorylation (A) and Src Y419 phosphorylation (B) after CSK incubation (10 min) followed by incubation with YopH to remove pY419 (1 h, 2 h, 3.5 h). Tyrosine phosphorylation of the Src C-terminus (A) was detected via dot blotting using Src pY530-specific primary antibody. Tyrosine phosphorylation of the Src activation loop (B) was detected via dot blotting using Src pY419-specific monoclonal primary antibody. (C) Time courses monitoring Src Y530 phosphorylation state of open (unphosphorylated) Src, Pyk2 and autoinhibited (pY530) Src, or Pyk2 and CSK. Each reaction includes 4 mM ATP. Tyrosine phosphorylation of the Src C-terminus was detected via dot blotting using Src pY530-specific primary antibody. (D) Time courses monitoring Src Y530 phosphorylation state for autoinhibited (pY530) Src Y419F with and without Pyk2. Tyrosine phosphorylation of the Src C-terminus was detected via dot blotting using Src pY530-specific primary antibody. *Inset:* Keys to immunoblotting antibody recognition sites in A, C, and D (Src pY530) and B (Src pY419), respectively.

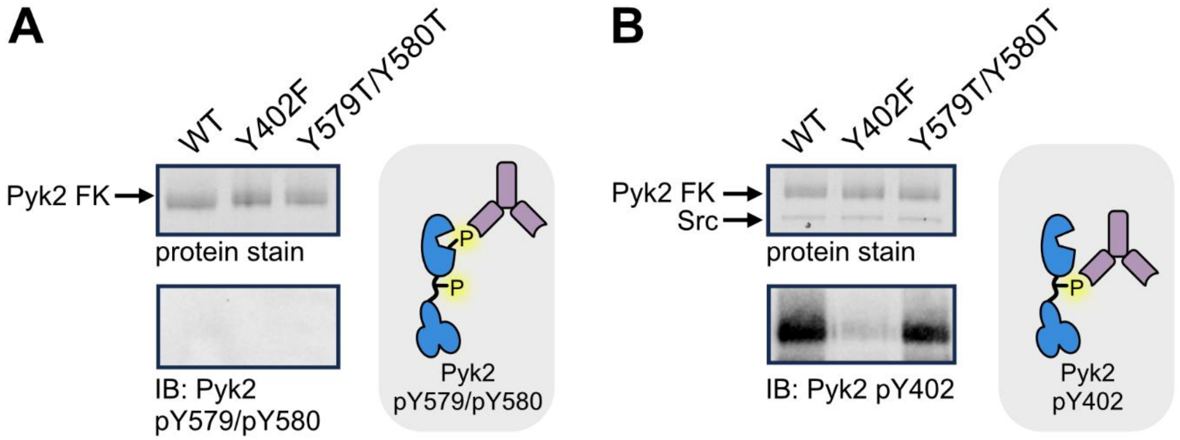


**Figure S6. Testing Pyk2 and Src phosphosite targeting specificity**
(A) Activation loop autophosphorylation of WT, Y402F, and Y579T/Y580T Pyk2 FERM–kinase (1 µM). Kinase variants were incubated with 4 mM ATP for 60 min at 37 °C. Phosphorylation of the Pyk2 activation loop was probed via Western blotting using Pyk2 pY579/pY580-specific primary antibody. (B) Phosphorylation of the FERM—kinase linker of WT, Y402F, and Y579T/Y580T Pyk2 FERM-kinase (1 µM) with basal state WT Src (0.2 µM) at 60 min. Tyrosine phosphorylation of the Pyk2 activation loop was probed via Western blotting using Pyk2 pY402-specific primary antibody. *Insets:* Keys to immunoblotting antibody recognition sites in A and B, respectively.

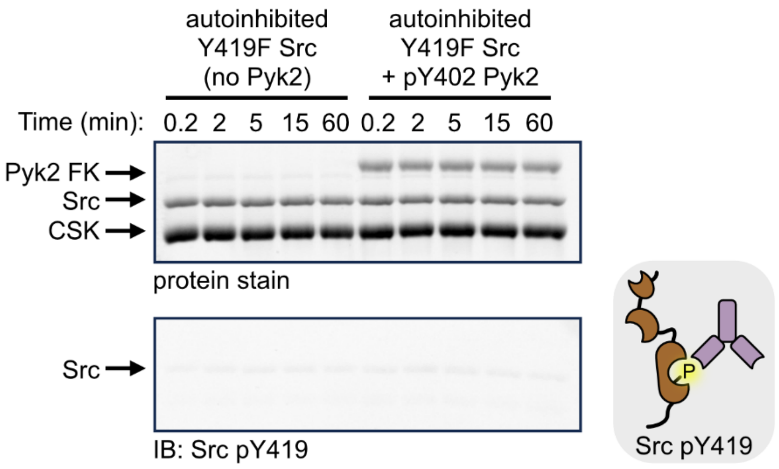


**Figure S7. Testing for autoinhibited Src phosphotransfer background**
To test for detectable phosphotransfer background of autoinhibited Src, pY530/Y419F Src (2 µM) was incubated in the presence or absence of pY402 Pyk2 FERM–kinase (2 µM) with 4 mM ATP. Autoinhibited, pY530/Y419F Src was generated as described in **Fig. 4**. Phosphorylation of the Src activation loop was probed via Western blotting using Src pY419-specific monoclonal primary antibody. *Inset:* Key to immunoblotting antibody recognition site.
